# Lineage-specific *tprK* diversification and *Treponema pallidum* transmission dynamics in Buenos Aires, Argentina

**DOI:** 10.64898/2026.01.29.702707

**Authors:** Nicole A.P. Lieberman, Luciana N. Garcia, Shah A. K. Mohamed Bakhash, Jeffrey C. Furlong, B. Ethan Nunley, Andrés Rabinovich, Patricia Fernandez Pardal, Viviana Leiro, Hong Xie, Farhang Aghakhanian, Mauro Romero Leal Passos, Wilma Nancy Campos Arze, Hugo Boechat Andrade, Silver K. Vargas, Kelika A. Konda, Michael Reyes Diaz, Carlos Caceres, Jeffrey D. Klausner, Jonathan B. Parr, Arlene Seña, Ariyaratne Manathunge, Lorenzo Giacani, Jaime Altcheh, Alexander L. Greninger

**Affiliations:** Department of Laboratory Medicine and Pathology, University of Washington School of Medicine, Seattle, WA, USA; Servicio Parasitología- Chagas, Hospital de Niños Ricardo Gutierrez, Capital Federal, Buenos Aires, Argentina; Instituto Multidisciplinario de Investigaciones en Patologías Pediátricas (IMIPP), CONICET, Buenos Aires, Argentina; Hospital de Infecciosas F J Muñiz, Servicio de Dermatología, Buenos Aires City, Argentina; Institute for Global Health and Infectious Diseases, University of North Carolina at Chapel Hill, Chapel Hill, NC, USA; Department of Microbiology and Parasitology, Biomedical Institute, Fluminense Federal University, Niterói, RJ, Brazil; Center for Interdisciplinary Studies in Sexuality, AIDS and Society, Universidad Peruana Cayetano Heredia, Lima, Peru; Division of Infectious Disease, David Geffen School of Medicine, University of California Los Angeles, Los Angeles, CA, USA; Keck School of Medicine, University of Southern California, Los Angeles, CA, USA; National STD/AIDS Control Programme, Ministry of Health, Colombo, Sri Lanka; Department of Medicine, Division of Allergy and Infectious Diseases, Harborview Medical Center, University of Washington, Seattle, WA, USA; Department of Global Health, Harborview Medical Center, University of Washington, Seattle, WA, USA; Vaccine and Infectious Disease Division, Fred Hutchinson Cancer Center, Seattle, WA, USA

## Abstract

**Background:** Syphilis rates are rising globally, with increases in congenital syphilis in South America particularly concerning. The characterization of contemporary South American *Treponema pallidum (Tp)* strains is crucial to syphilis vaccine development, yet few genomic epidemiology studies have focused on this region. Here, we performed whole genome sequencing (WGS) of *Tp* from Buenos Aires, Argentina, as well as deep sequencing of the hypervariable *tprK* locus, which is critical to *Tp* immune evasion.

**Methods:** People with primary, secondary, or congenital syphilis were enrolled at two clinics in Buenos Aires between October 2018 and January 2023, including individuals associated with intra-household transmission. Hybrid capture WGS was performed and a core genome phylogeny generated. K-mer-based methods using full-length *tprK* PacBio long reads were used to uncover differences in diversity and detect *Tp* transmission.

**Findings:** *Tp* genomes were recovered from 70 individuals in Buenos Aires and primarily belonged to globally dominant SS14 sublineage-1 and Nichols sublineage-8, as did *Tp* from Brazil (n=8). Peruvian samples (n=3) all belonged to sublineage-1. Two individuals from Argentina had co-infections with Nichols- and SS14-lineage strains. Macrolide resistance via A2058G occurred in 27/70 (38.6%) samples. Across 56 samples, *tprK* allelic diversity was significantly increased in secondary syphilis, oral lesions, and SS14-lineage strains compared to primary syphilis, anogenital lesions, and Nichols-lineage strains, respectively. Increased diversity in SS14-lineage strains is driven by an enhanced repertoire of V7-specific donor sequences. *tprK* sequences from intra-household transmissions were more similar than unrelated samples with identical core genomes.

**Interpretation:** *Tp* circulating in South America is closely related to dominant global sublineages. Increased *tprK* diversity in the SS14 lineage may influence *Tp’s* ability to escape host immunity. *tprK* profiling is a promising tool to elucidate syphilis transmission networks. This study underscores the utility of genomics to yield insights into *Tp* pathogenesis.

**Research in context:** *Evidence before this study:* Although whole genome sequencing (WGS) and genomic epidemiology have contributed to an understanding of the global diversity of *Treponema pallidum* (*Tp*), very few strains from South America have been sequenced to date. On January 6^th^, 2026, we performed a PubMed search including terms “syphilis genomic epidemiology”, “South America”, “*Treponema pallidum*”, and “Argentina”. We excluded studies of ancient *Tp* samples found in South American archeological sites. A single *Tp* sample from Argentina was originally sequenced in 2016 and included in subsequent analyses of global diversity. Nine Peruvian samples were previously sequenced in a study of global diversity on six continents. Thirty-three samples from Cali, Colombia were predominantly SS14-lineage and genotypic macrolide resistance was found in half of strains. Additionally, studies of *Tp* in Buenos Aires using a multi-locus sequence typing approach have shown circulation of strains belonging to both Nichols and SS14 lineages, with an increasing rate of macrolide resistance over time. Five previous studies have examined *tprK* in clinical specimens and consistently shown increased *tprK* diversity in specimens associated with secondary syphilis compared to primary syphilis lesions. One study has shown that loss of *tprK* donor cassettes is associated with reduced *tprK* diversity, consistent with results from *in vitro* experiments. A single study has shown that *tprK* sequence content is more similar in samples with a suspected epidemiologic link, though no prior studies have looked at *tprK* diversity in the context of known syphilis transmission events.

*Added value of this study:* This study is the first to use WGS to comprehensively examine *Tp* transmission in a large South American city, yielding 96 samples from 70 individuals. We also add the first contemporary *Tp* genomes from Brazil. In contrast to findings from England, Australia, and other high-income countries, no *Tp* sublineages are associated with demographic groups or sexual networks in Buenos Aires. Two of seventy patients were co-infected with both Nichols- and SS14-lineage strains, showing the previously unappreciated frequency of conditions that permit inter-strain recombination-driven diversification of *Tp*. This study also reveals novel aspects of *Tp* pathogenesis, including lineage-specific differences in *tprK* diversity. We also develop methods for the analysis of *tprK* relatedness between samples and demonstrate that *tprK* sequences are more similar in samples from individuals within intra-household syphilis transmission chains compared to those from epidemiologically unrelated individuals.

*Implications of all the available evidence:* *Tp* strains circulating in Buenos Aires are genetically similar to those circulating worldwide and in Brazil and Peru but are noteworthy for the low (but rising) rate of macrolide resistance. Lineage-specific patterns of *tprK* antigenic variation could result in differences between Nichols- and SS14-lineage strains’ interactions with the host immune system. Finally, we show that *tprK* profiling holds promise to identify samples from within a syphilis transmission chain and could play an important role in public health.

## Introduction

In 2022, the Region of the Americas accounted for 42% (3.37 million) of all new syphilis diagnoses worldwide, representing the highest incidence of any World Health Organization region. Low- and middle-income countries in Central and South America account for most cases. The rate of syphilis in pregnant women in the Region increased 28% between 2020 and 2022, with 68,000 cases of congenital syphilis annually^1^ and in Argentina in 2023, 6% of women attending antenatal care tested positive for syphilis^2^. Until a vaccine that can prevent vertical transmission of *Treponema pallidum (Tp)* is available, reducing rates of congenital syphilis depends on identifying and breaking transmission chains, a process complicated by overlapping sexual networks and high rates of syphilis in vulnerable and hard-to-reach populations^3,4^. Despite the key role genomic epidemiology can play in understanding the spread and evolution of *Tp*, South American strains have been the subject of limited study.

Genomic studies of clinical samples can also shed light on the basic biology of *Tp*, which is difficult to study in the laboratory due to technically challenging cultivation and limited genetic tools^5,6^. *Tp* is notable for its slow accumulation of genetic diversity, with fewer than 150 single nucleotide variants (SNVs) in the core genome separating strains of the Nichols and SS14 lineages of *Tp*^7,8^, which are clinically indistinguishable^9^. In contrast, ongoing gene conversion in *tprK* results in antigenic variation (AV) of seven extracellular loops of the outer membrane protein TprK. Donor cassettes (DCs) from 53 chromosomal sites are recombined into the seven variable (V) regions to generate novel *tprK* alleles, tens or hundreds of which may be found in primary or secondary syphilis lesions^10,11^, enabling evasion of host adaptive immune responses^12,13^. Understanding TprK AV can provide insight into *Tp*-immune dynamics that cause the relapsing clinical course of syphilis^13^ and *tprK’*s rapid evolution may be used to identify syphilis transmission networks^14^ where whole genome sequencing (WGS)-based methods lack resolution due to low diversity of the *Tp* core genome.

Here, we use WGS to characterize contemporary *Tp* in Argentina, Brazil, and Peru. In the Argentinian cohort, we also investigate *tprK* sequence diversity by *Tp* lineage and leverage previously-identified cases of intra-household syphilis transmission^15^ to refine our *tprK-*based methods to deconvolute syphilis transmission networks^14^.

## Methods

### Ethics Approvals

In Buenos Aires, enrollment of pediatric participants and their caregivers (DI-2018-245-HGNRG) as well of adult participants (DI-2022-155-HGNRG) received approval from the Committee and Review Board of the Hospital de Niños Ricardo Gutierrez. Sequencing of *Tp* from deidentified specimens was approved by the University of Washington IRB (STUDY00000885) and the University of North Carolina at Chapel Hill IRB (24-2696). Written consent was obtained from all adult participants and caregivers of enrolled children.

### Enrollment Criteria and Screening

A convenience sampling method was used to recruit patients in two public health services in Buenos Aires, Argentina between October 2018 and January 2023. A total of 94 adult outpatients were assisted in the Dermatology Unit of the Hospital Javier Muñiz, whereas another cohort of 47 patients, including pediatric outpatients (n=28) and their caregivers (n=19) were assisted at the Hospital de Niños Ricardo Gutierrez as previously described^15^. Pediatric cases where sexual abuse was suspected were excluded. Additional screening criteria is found in Supplementary Methods, as are enrollment criteria for Brazil and Peru cohorts, which were included in WGS analyses.

### *T. pallidum* WGS, genome assembly, and phylogenetics

WGS was performed as previously described^16^. Ninety-six samples from 70 participants from Argentina, 8 samples from 8 participants from Brazil and 3 samples from 3 participants from Peru were newly sequenced for this study and included in a phylogenetic analysis alongside 1,772 publicly available genome sequences from the NCBI Sequence Read Archive (Supplementary Data). Genomes were assembled and variants called using a custom pipeline available at https://github.com/greninger-lab/tpallidum-variant-calling, outlined in Supplementary Methods.

### *tprK* sequencing and data pre-processing

The *tprK* gene was amplified as previously described^16^ in technical duplicate and sequencing performed on the PacBio Revio. Following demultiplexing, HiFi reads were filtered to remove sequences shorter than 1400 bp or longer than 1700 bp or with unexpected primer orientations. Only reads with each base Q30 or higher were included in subsequent analyses. Variable regions and intervening constant regions were translated and reads containing frameshifts or premature stop codons in any region excluded from analysis. Custom analyses are described in Supplementary Methods.

### Statistics and Reproducibility

All calculations and statistics were performed in R v4.2.2. Custom processing scripts are available at https://github.com/greninger-lab/tprk-argentina/.

### Data Availability

All WGS data generated during this study are available at NCBI BioProject PRJNA723099, with accessions in Supplementary Data. *tprK* PacBio raw data are available in NCBI BioProject PRJNA1415869.

## Results

### Cohort demographics

A total of 141 individuals with syphilis (113 adults and 28 children) were enrolled, including seven individuals associated with three intra-household transmissions. Demographic characteristics of the adult and pediatric cohorts are summarized in Table 1. All adults were cisgender, with median age of 26 years (IQR 22-33). All 45 men who have sex with men (MSM) were adult males, as were all 27 persons living with HIV (PLWH), including five individuals newly diagnosed during enrollment. Twenty-eight patients had prior syphilis. The pediatric cohort included four newborns with congenital syphilis, 12 children (five males and seven females) with a median age of 2 (IQR: 1-6) years and 12 adolescents (seven males and five females) with a median age of 16 (IQR: 15-17) years. Adolescents with primary and secondary syphilis reported sexual activity. Children with secondary syphilis caused by non-sexual close contact with active lesions from their caregivers^15,17^ have a median age of 3 years (IQRs: 2-6). Clinical findings are included in Supplementary Analysis.

**Table 1.** Demographic characteristics by cohort. *The p value was derived from a chi-square test comparing the proportion of patients with sequenced genomes to the total patient population.

|  | Adult<br>patients<br>(n=113)<br>n (%) | Pediatric<br>patients<br>(n=28)<br>n (%) | Total Patients<br>(n=141)<br>n (%) | Patients with<br>WGS<br>(n=70)<br>n (%) | p<br>value<br>* |
| --- | --- | --- | --- | --- | --- |
| AGE: median<br>(IQRs) years | 26 (22-33) | 6 (2-16) | 24 (19-31) | 27 (18-33.5) |  |
| Adults (>18) | 113 (100.0) | - | 113 (80.1) | 51 (72.8) | 0.31 |
| Pediatric (0-18) | - | 28 (100.0) | 28 (19.9) | 19 (27.2) |  |
| **BIOLOGICAL SEX** |  |  |  |  |  |
| Male | 78 (69.0) | 13 (46.4) | 91 (64.5) | 48 (68.6) | 0.67 |
| Female | 35 (31.0) | 15 (53.6) | 50 (35.5) | 22 (31.4) |  |
| **SEXUAL ORIENTATION (among adult and adolescent males)** |  |  |  |  |  |
| MSM | 45 (57.7) | 0 (0) | 45 (52.9) | 19 (43.2) | 0.39 |
| No MSM | 33 (42.3) | 7 (100%) | 40 (47.1) | 25 (56.8) |  |
| **HIV** |  |  |  |  |  |
| Positive | 26 (23.0) | 0 (0) | 26 (18.4) | 9 (13.8) | 0.41 |
| Negative | 87 (77.0) | 28 (100%) | 115 (81.6) | 61 (86.2) |  |
| **PRIOR SYPHILIS** |  |  |  |  |  |
| Yes | 27 (23.9) | 1 (3.6) | 28 (19.8) | 10 (14.2) | 0.28 |
| No | 85 (75.2) | 27 (96.4) | 112 (79.4) | 70 (85.8) |  |
| Not specified | 1 (0.9) | 0 (0) | 1 (0.7) | 0 (0) |  |
| **CLINICAL STAGE** |  |  |  |  |  |
| Primary | 21 (18.6) | 3 (10.7) | 24 (17.0) | 16 (22.8) | 0.24 |
| Secondary | 92 (81.4) | 21 (75.0) | 113 (80.1) | 54 (77.2) |  |
| Congenital syphilis | 0 (0) | 4 (14.3) | 4 (2.8) | 0 (0) |  |

### Yield of molecular testing and WGS

A total of 250 samples were collected from the 141 enrolled individuals, primarily oral and anogenital swabs (135/250, 54.0% and 92/250, 36.8% of samples, respectively). Molecular positivity for the single copy *tp0574* locus was highest among anogenital swabs (81/92, 88.0%), extragenital rash (9/13, 69.2%), and oral swabs (81/135, 60%). All 22 primary chancre swabs were positive. Two of thirteen (15.4%) blind swabs of normal mucosa in primary syphilis were *tp0547* positive. Among secondary syphilis, swabs of oral lesions were positive in 37 of 42 cases (88.1%), while blind swabs of normal oral mucosa were positive in 36 of 66 cases (54.5%) (Figure S1A). Primary lesions had the highest interpolated *tp0574* copies per swab (median: 48,121 copies, n=16), followed by secondary disseminated anogenital lesions (median: 17,025 copies n=39) (Figure S1B). Specimens from individuals with prior episodes of syphilis (n=44 specimens from 28 individuals) were less frequently *tp0574* positive than those without (23/44, 52.3% vs 153/204, 75.0%), p=0.0047, chi-square test).

We attempted WGS on samples for which we could input >200 genome copies to library preparation. Ninety-six samples from 70 patients had sufficient coverage for analysis, which we defined as having less than 25% ambiguous bases in SNV positions. The highest yield of near-complete genomes was from primary anogenital lesions (16/22, 72.7%; Figure S1C), followed by oral and anogenital mucosal lesions. No near-complete genomes were recovered from blood. The proportion of genomes recovered among demographic groups was representative of the enrolled cohort (Table 1, all p>0.05), although no genomes were recovered from infants with congenital syphilis.

### Both lineages of *T. pallidum* circulate in Buenos Aires

Prior studies have shown that SS14- and Nichols-lineage strains circulate in Buenos Aires^15,18^ and Colombia^9,19^. We built upon these studies by analyzing the genomes recovered from Argentine samples alongside publicly available South American *Tp* sequence data as well as three new samples from Peru and eight from Brazil (Supplementary Methods). First, we contextualized South American strains alongside a total of 1,879 *Tp* samples (Figure 1A). Most South American samples are closely related (within 10 SNVs) to previously classified samples in globally distributed sublineages 1 (SS14 lineage) and 8 (Nichols lineage)^8^ (Figure 1B). In Argentina, 32/71 (45.1%) individuals had Nichols-lineage treponemes, similar to Colombia (8/23, 34.5%). Although genotypic macrolide resistance, conferred by A2058G mutation in the 23S rRNA gene, was relatively low in Argentina compared to other South American countries (39.1%; Figure 1C), these data reflect a progressive increase in resistance within Buenos Aires over the past decade^15,18,20^.

**Figure 1:**
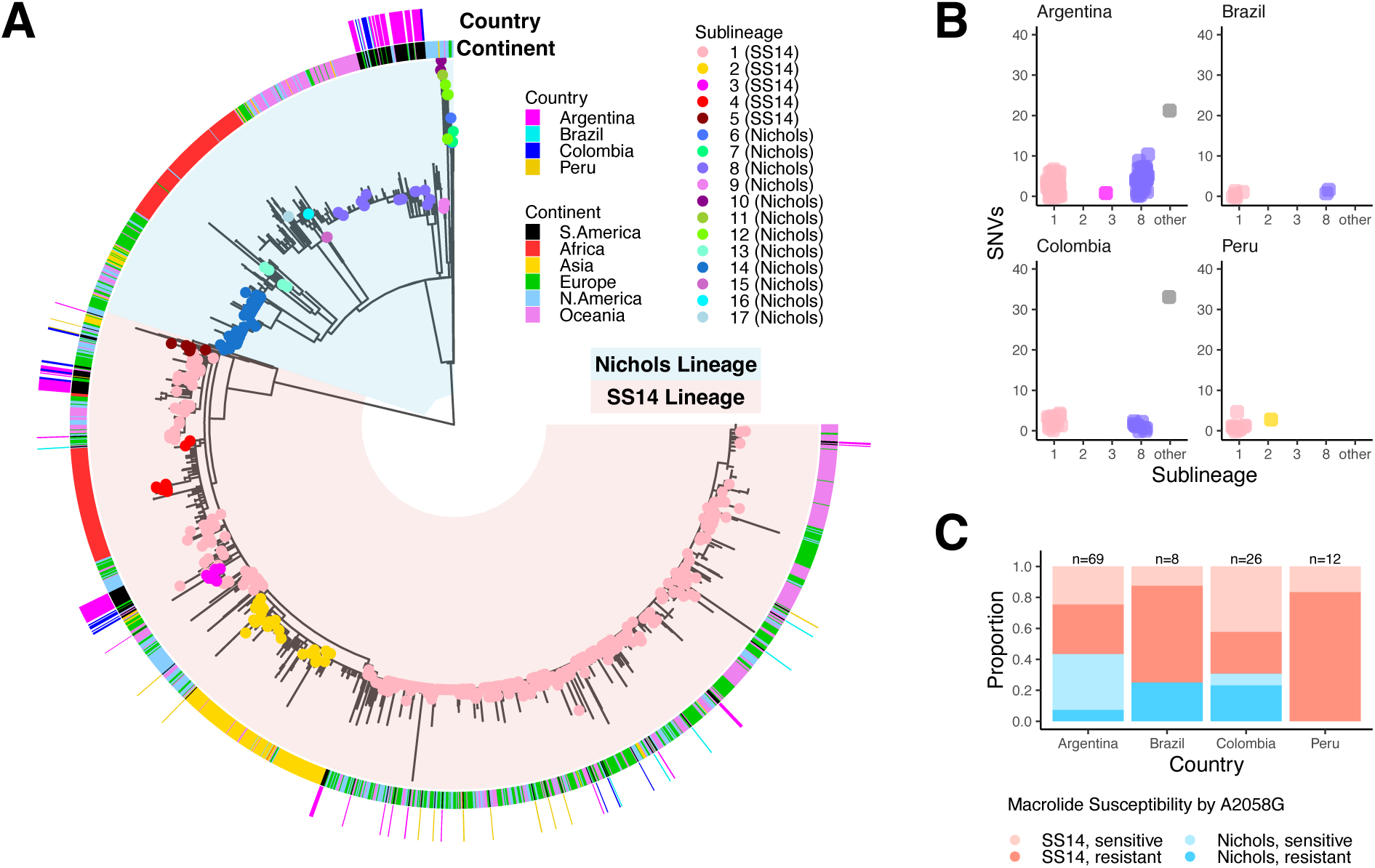
South American *T. pallidum* is closely related to globally dominant sublineages. A) Recombination-masked, maximum likelihood phylogeny of 1,879 *T. pallidum* strains, including 107 newly sequenced for this study. Blue background represents the Nichols lineage and red represents the SS14 lineage. Strains included in a study by Beale *et al* (2021) have tips colored by sublineage. The inner ring denotes continent of collection and outer ring denotes country for South American samples only. B) Number of single nucleotide differences between each South American sample and its most closely related worldwide sample, plotted by sublineage of the worldwide sample. C) Presence of A2058G mutation conferring macrolide susceptibility, by country and lineage.

Two individuals in the Buenos Aires cohort had evidence for co-infection, reported only once previously. In one individual, mixed sequence reads at lineage defining positions^8^ were detected in a residual genital chancre (Figure 2A). In another person, SS14- and Nichols-lineage strains were recovered from different lesions (Figure 2B). Both individuals reported multiple sex partners in the preceding three months. Co-infections were confirmed by strain-specific digital PCR^21^ using DNA re-extracted from the original swab. In one of 20 individuals with WGS from multiple secondary syphilis lesions, we found compartmental SNVs in *tp0006*, a gene of unknown function (Figure S2, Supplementary Analysis).

**Figure 2:**
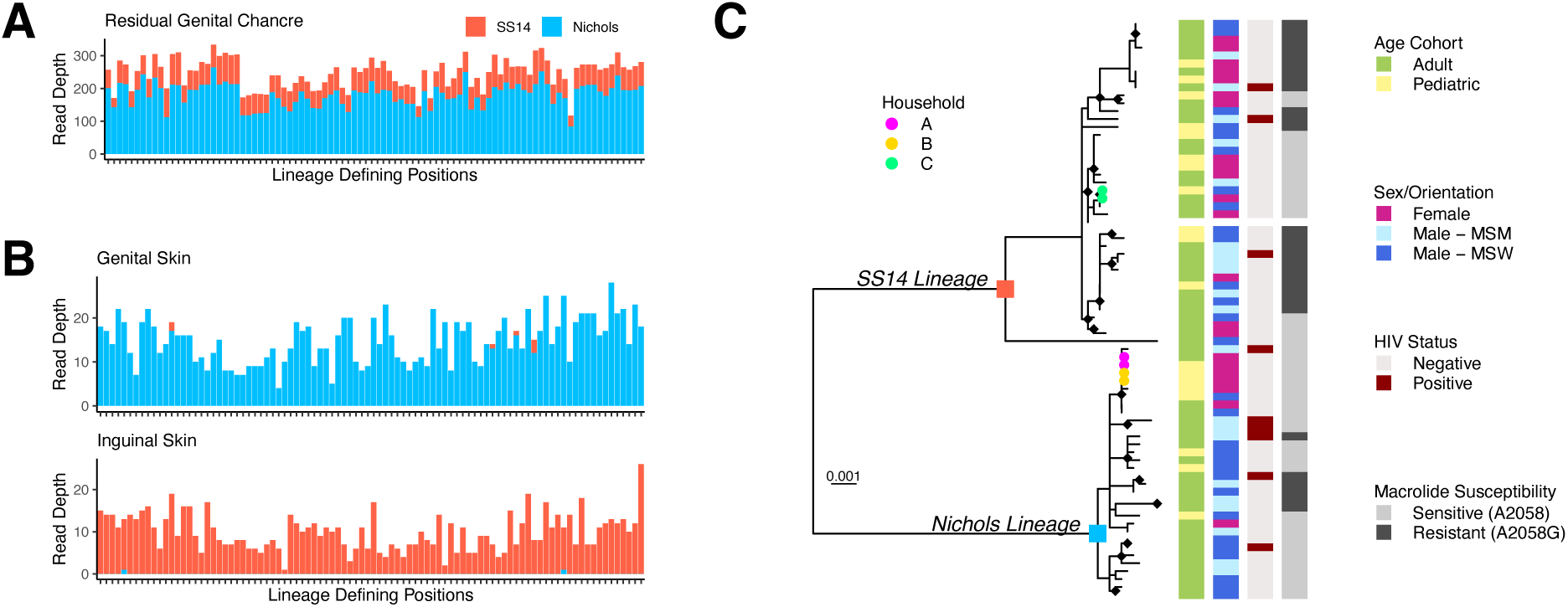
Co-infection and *Treponema pallidum* genomic epidemiology within Buenos Aires, Argentina, 2018-2023. A) Whole genome sequencing read depth shows the presence of ∼80% Nichols- and ∼20% SS14- variants at lineage-defining positions, supporting a co-infection in the residual genital chancre of an individual (GSPAR041) with secondary syphilis. B) Secondary syphilis lesions from genital and inguinal skin of another individual (GSPAR083) belong to Nichols and SS14 clades, respectively. Individuals in both A and B had a history of multiple sex partners. C) Phylogeny of strains collected in Buenos Aires, 2018-2023. Nichols and SS14 lineage ancestral nodes are labeled with blue and red squares, respectively. Tips from samples of each patient within the three household transmission clusters are colored red, green, and blue, respectively. Tiles correspond to patient age, sex and sexual orientation, HIV status, and macrolide susceptibility (A2058G).

### No association between sublineage and demographic characteristics in Buenos Aires

We constructed a phylogeny of *Tp* genomes from Buenos Aires, including strains from our current study and one sample collected in 2016^15^ (n=73 genomes from 71 individuals, Figure 2C). Thirty-one samples formed nine distinct clusters with 0 core-genome SNVs, including clusters of individuals within households as well as clusters with no known epidemiologic link. Among individuals without Nichols-SS14 co-infection and with clinical information (n=68), we did not find any association between patient sex, sexual orientation, or HIV status with *Tp l*ineage. Macrolide resistance was more prevalent in the SS14 lineage than the Nichols (22/38, 57.9% vs 5/30, 16.7%, p<0.01), and, among adults and adolescents, in MSM (12/17, 70.5%) versus MSW or females (10/25, 40% (p=0.05) and 5/17, 29.4% (p<0.01), respectively). Mutations to 16S rRNA sites homologous to those conferring tetracycline resistance were not observed^22,23^.

### The number and distribution of *tprK* alleles depends on stage, site, and *T. pallidum* lineage

We next interrogated the hypervariable *tprK* gene using high fidelity PacBio sequencing coupled with our customized analyses (**Figure S3**). TprK undergoes antigenic variation within seven of its extracellular loops to evade host adaptive immunity^12,13,24–26^ by replacing fragments of *tprK* with donor cassettes (DCs) from 53 chromosomal regions arranged in two loci (Figure S4). Variable regions were defined essentially as described previously^10,24^, however, for ease of interpretation we split variable and constant regions between codons. Sequences between variable loops exhibited minimal variation, with no changes to a known B cell epitope^27^ (Figure S5, Supplementary Analysis).

We examined the relationship between clinical and demographic factors in the number of unique *tprK* alleles, V regions, and Shannon diversity index in each sample, including a total of 56 samples from 44 individuals with sufficient read depth and quality (Figure S3). V regions were only counted if present in both technical replicates. We did not limit inclusion of full-length *tprK* alleles to only those present in both technical replicates because it had a different effect on primary and secondary samples (Figure S6A-B), consistent with differences in the underlying distribution of alleles between primary and secondary syphilis^11^. No full-length *tprK* alleles recovered were shared between patients. Among ten individuals with *tprK* sequencing from more than one body site, six paired samples shared one or two full-length *tprK* alleles. The number of *tprK* alleles exceeding 0.2% abundance ranged between 7-141 per sample. Consistent with prior reports^10^, secondary syphilis samples of all types yielded a significantly higher number of unique full-length *tprK* alleles (median 63 vs 37, p <0.001, Figure 3A) as well as more V regions and higher Shannon diversity index. When lesion type and location was considered, oral secondary lesions had significantly higher *tprK* diversity than anogenital primary lesions, though all secondary lesion types except rash had a higher median number of alleles and greater diversity than primary lesions (Figure 3B). Among the five individuals from whom paired oral and anogenital lesions of any type were collected, oral lesions consistently had higher *tprK* diversity than anogenital lesions (Figure 3C). Among 55 samples from 43 patients with paired WGS and full-length *tprK* profiling, SS14-lineage samples had more unique *tprK* alleles than those from the Nichols lineage (Figure 3D), driven primarily by increased diversity in V6 and V7 (Figure 3E, Figure S7). Among samples with paired WGS, a total of 3,142 unique *tprK* alleles were found, including 1,191 in Nichols-lineage samples (n=25) and 1,951 in SS14-lineage samples (n=30). Sex, sexual orientation, prior syphilis, HIV coinfection, and age were not associated with differences in *tprK* diversity (Figure S8).

**Figure 3:**
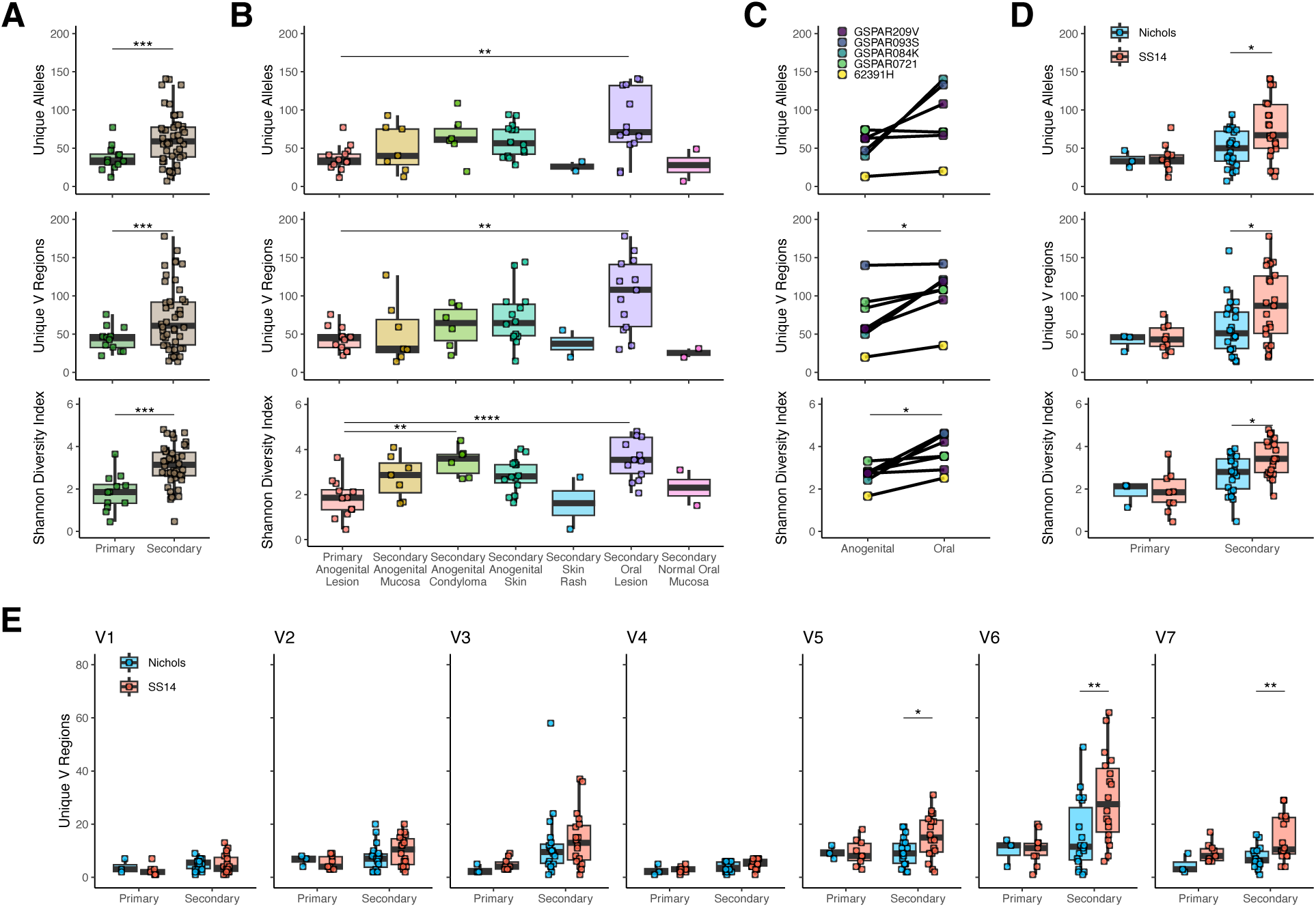
Number and distribution of *tprK* alleles. A) Differences in total alleles (top), unique variable regions (middle), and Shannon diversity index (bottom) by syphilis stage. T-test, ***p<0.001. B) Differences in *tprK* by lesion type and location. ANOVA with Tukey’s post hoc, **p<0.01, ****p<0.0001. C) Samples from individuals with paired samples from anogenital and oral sites show a higher number of V regions and diversity index in the oral lesion. Paired T-test, *p<0.05. D) *tprK* diversity in Nichols vs SS14 lineage samples, t-test, *p<0.05. E) Variation by clinical stage and lineage in the number of unique V regions. T-test, *p<0.05, **p<0.01.

### SS14 lineage strains generate *tprK* diversity with an expanded repertoire of *tprK* donor cassettes

To further explore the observation of increased *tprK* diversity in the SS14 lineage, we applied a k-mer-based analytical approach designed to account for its extensive rearrangement. *tprK* evolves via recombinogenic changes that replace segments of tens of nucleotides in each variable region^24^ (Figure S3B), precluding the use of maximum-likelihood methods to determine sequence relatedness. Therefore, we examined differences in *tprK* sequence composition between lineages by generating overlapping subsequences (k-mers) of lengths ranging between 9-27nt (Figure S9), allowing comparison of sequence similarity despite rearrangement. We focused particularly on k-mers of length 21 nt since the seven-amino acid fragments (7AA-mers) they encode approximate the minimal length of a linear B cell epitope^28^, yet are short enough to enable identification of *tprK* sequence fragments that are exact matches to chromosomal donor cassettes (DCs)^24,29^. We defined *tprK* 21-mers that perfectly matched one or more DCs as “DC-derived”, in contrast to mosaic 21-mers that were stitched together from multiple DCs. K-mers and AA-mers found in *tprK* sequences from strains of only the Nichols or SS14 lineage are defined as “lineage-specific”.

To ensure balanced representation of 21-mer diversity within each clade, SS14 samples (n=30) were first down-sampled to the same number (25) as Nichols, then 1,000 *tprK* alleles from each lineage randomly selected from the 25 samples. SS14-lineage samples had more V6 and V7 21-mers (and 7AA-mers) than Nichols (Figure 4A-B), with a considerably greater proportion of unique SS14 7AA sequences derived from V7 than in Nichols (Figure 4C). While SS14 and Nichols strains had similar proportions of total DC-derived 7AA fragments, SS14 strains had significantly more lineage-specific DC-derived 7AA fragments than Nichols (44 vs 9, p<0.01), half of which were found in V7 (Figure 4D). This suggests the higher diversity in SS14 lineage *tprK* can be at least partially attributed to cassettes of 21 nt or longer being inserted from a donor locus directly, as opposed to shorter fragments being stitched together to create mosaics not found in the donor loci.

**Figure 4:**
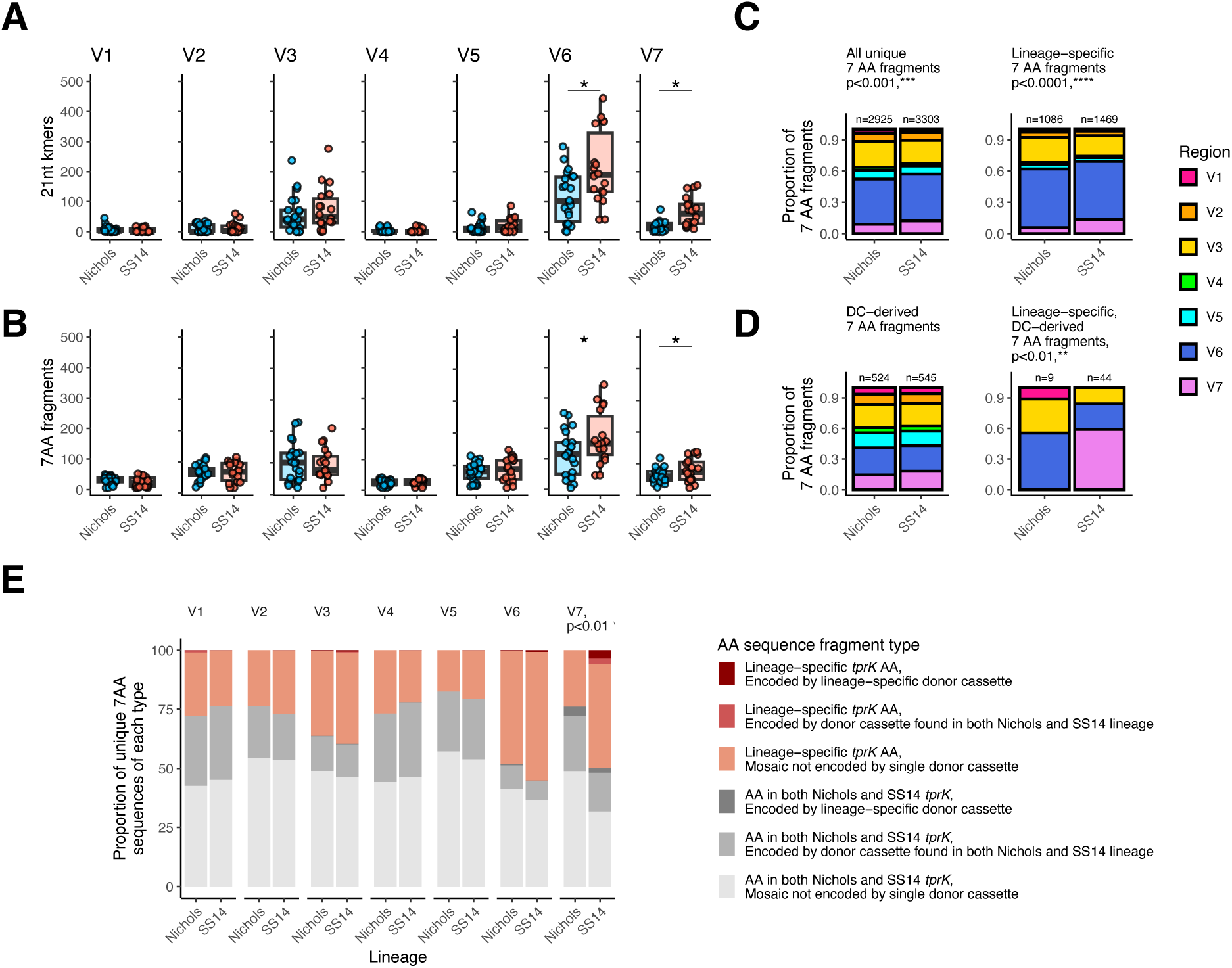
Lineage-specific differences in *tprK* sequence content. All analyses in this figure were performed by sampling an equal number of samples from each lineage (25) followed by random subsampling of 1,000 unique alleles per lineage. Significance is defined as a mean p<0.05 across 1,000 independent subsampled replicates. A single representative replicate is visualized. The total number of unique V6 and V7 21 nt k-mers (A) and 7AA-mers (B) are significantly higher in SS14 than Nichols lineage. T-test, *p<0.05. C) The proportion of unique 7AA-mers derived from each region differs by lineage among all 7AA-mers as well as those found in only the SS14 or Nichols lineage. ***p<0.001 and ****p<0.0001, chi squared. D) The proportion of 7AA-mers derived from each region and encoded directly by donor cassettes is not different between SS14 and Nichols lineages, but each lineage contains a different proportion of lineage-specific 7AA-mer fragments directly encoded by donor cassettes, **p<0.01, chi squared. SS14 uses more lineage-specific donor cassette (DC)-derived sequences than Nichols (44 vs 9). E) Proportion of unique 7AA-mers of each type – present in both lineages, Nichols only, or SS14 only, and encoded by a single DC or a mosaic sequence not found in the donor cassettes – by variable region. SS14 lineage strains contain significantly more lineage-specific V7 7AA-mers than Nichols, including both DC-derived and mosaic sequences generated by gene conversion, **p<0.01, chi squared.

We previously showed how differences in donor repertoire can affect *tprK* diversity by genetically knocking out the major DC locus^26^ and detecting a clinical strain missing nine sets of DCs that had a reduced number of *tprK* variable regions relative to wild type strains^14^. Among strains in this analysis, differences between Nichols and SS14 DC loci were confined to SNVs in DCs specific for V3, V6, and V7, as well as SNVs between DCs (Figure S10). The majority of lineage-specific DC-derived 21-mers contained those lineage-specific SNVs, and most strains harbored at least one *tprK* allele with a lineage-specific 7AA-mer derived from the V3/V6 overlapping DC sites 26 and 25 (Figure S11A). The largest difference between Nichols and SS14 DCs is a 51bp indel in DC site V7-31, which directly contributes novel sequences to *tprK* alleles in SS14-lineage samples. When assessing both the type (DC-derived or not) and lineage specificity of unique *tprK* k-mers, we showed that SS14 has significantly greater use of lineage-specific V7 7AA fragments than Nichols (Figure 4E). Interestingly, the lineage-specific DC sequences found in Nichols V7-31 and V7-2 are not found in Nichols *tprK* alleles, while SS14-specific sequences are readily found in SS14 alleles (Figure S11A-B), thus accounting for the reduced diversity of the V7 *tprK* allele seen in Nichols strains. Together, these data suggest that the increased *tprK* sequence diversity observed in SS14-lineage strains is due in part to an enhanced repertoire of V7 donor cassettes.

### *tprK* is more similar between individuals within known transmission chains than between unrelated individuals

We used our prior methods to confirm and expand our finding that *tprK* sequences shared by two individuals are more similar in people with a suspected epidemiologic link^14^. The inter-sample Pearson correlation between the proportion of V regions shared between samples is greater within known intra-household transmission events than in epidemiologically unrelated samples with identical core genomes (Figure 5A, p<0.05, t-test)^14^. Next, we increased the specificity of the comparison by determining the inter-sample Bray-Curtis dissimilarity for each V region and length of k (Supplementary Methods). Each variable region had a differential ability to discriminate between samples from the same patient, same household, or unrelated individuals (Figure S12). V1 and V4 have lower Bray-Curtis dissimilarity between samples from the same household than unrelated individuals, while V6 only has low k-mer dissimilarity in samples from the same individual. These results are consistent with prior observations that V1 and V4 evolve the most slowly and V6 evolves the most rapidly, resulting in the highest diversity among V regions^29^. We calculated a composite Bray-Curtis dissimilarity score that included k-mers from all seven V regions, summed across all values of *k* (Figure 5B) as well as the median inter-allele global alignment distance between *tprK* alleles within and between each sample (Figure 5C). Finally, we determined the median inter-allele composite Bray-Curtis score (Figure 5D). All methods showed a significant difference in *tprK* similarity between known transmission chains (intra-household) versus epidemiologically unrelated samples. Analysis of full-length alleles (Figure 5C-D) did not further resolve distinctions between known transmission events and unrelated samples relative to methods relying only on total V region content (Figure 5A-B).

**Figure 5:**
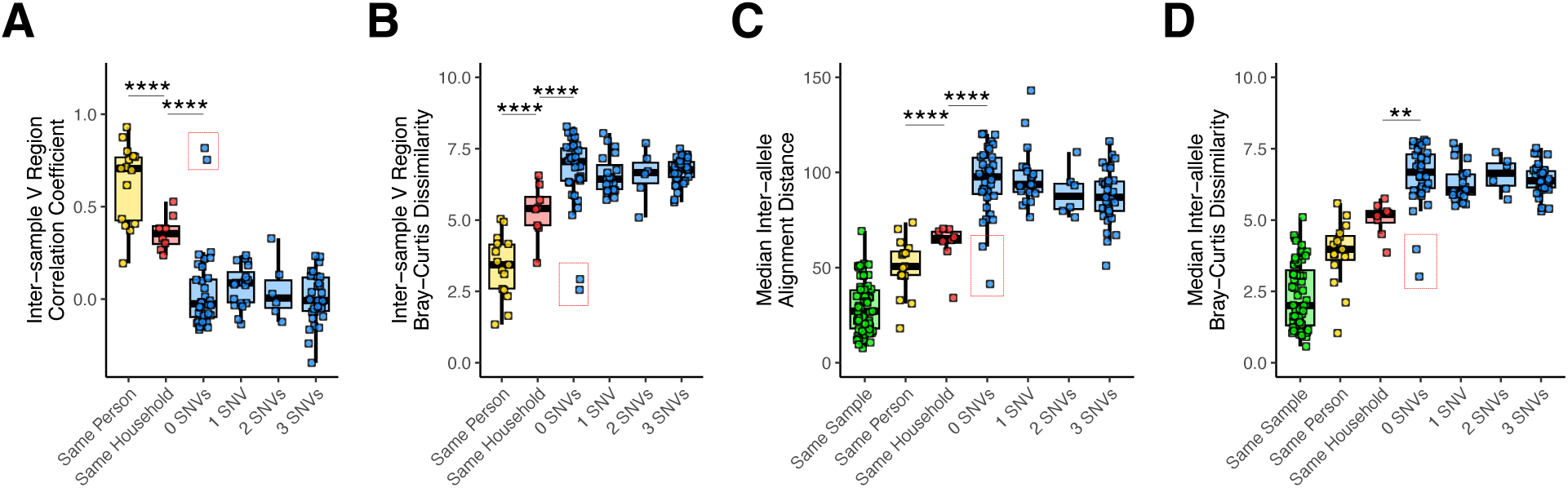
Inter-sample *tprK* relatedness by pairwise sample relationship. Pairwise comparisons of *tprK* profiling data were performed using different statistical methods. Samples were grouped based on whether they were taken from different samples from the same individual, individuals from the same household constituting likely transmission, and based on core genome SNV distance. A) Pearson coefficient of relative proportions of variable regions. B) Composite Bray-Curtis Dissimilarity for DNA kmers in variable regions. C) Median pairwise alignment distance between *tprK* alleles in each sample pair type. D) Median composite inter-allele composite Bray-Curtis dissimilarity in each sample pair type. ANOVA with Tukey’s post hoc, **p<0.01, ****p<0.0001. Red boxes enclose samples from a pair of individuals that may be epidemiologically linked based on patient age, sex and orientation, and presentation to the clinic on the same day,

Interestingly, among samples separated by 0 SNVs, all methods identified unexpected similarity between the oral and genital samples from one patient and the genital sample from a second patient (Figure 5, red boxes). Notably, outside of the V regions, strains from both patients also had a W516C mutation in the C-terminus of TprK (Figure S5B). Upon further review of patient clinical data, we learned that the samples came from a male with secondary syphilis and a female with primary syphilis. Both patients were in their mid-20s, heterosexual, denied stable sexual partners, and visited the clinic on the same day. Together, these findings support the utility of *tprK* sequencing in identifying and prioritizing syphilis transmission chains for epidemiological investigation.

## Discussion

As the syphilis epidemic in South America accelerates, sequencing of *Tp* provides considerable insight into syphilis epidemiology, revealing that most strains isolated in South America are closely related to those circulating throughout the world^8^. We also extend our previous methods using *tprK* similarity to elucidate transmission events, which can augment public health responses by detecting network links not captured by traditional contact tracing approaches^3^. Using multiple independent methods to profile *tprK* relatedness in a unique cohort of samples that included household linkage, we show *tprK* sequences from individuals within a transmission chain are more closely related than those from unrelated samples with identical core genomes. Moreover, the method revealed a possible transmission event that had not been recorded clinically, showing the power of this technique to identify closely related cases that merit follow-up public health investigation. Analysis of full-length *tprK* alleles did not demonstrate a significantly enhanced ability to resolve inter-sample distances relative to analysis of V regions alone, suggesting that *tprK* profiling could be deployed using short-read sequencing technologies that have been widely adopted in public health laboratories.

An important limitation of the current study of *tprK* is the use of convenience sampling from a single location, Buenos Aires, resulting in a relatively small number of participants for a very diverse metric like *tprK*. In particular, the number of known transmission events (three) was very low. Furthermore, the household transmission samples – suspected to be acquired syphilis by nonsexual contact^15,17^ - may not be fully representative of *tprK* evolution during sexual transmission. Nonetheless, the work described here replicates and significantly extends prior *tprK* profiling in Seattle, Washington^14^.

In addition to the key public health questions that can be answered using *Tp* WGS and targeted *tprK* sequencing, genomic studies of *Tp* evolution in patient samples can reveal aspects of pathogenesis that are challenging to study in vitro given limited experimental tractability and technically challenging cultivation. Unexpectedly, we showed that *tprK* sequences recovered from oral lesions are more diverse than from anogenital lesions, even from patient-matched specimens, which may reflect different immune surveillance or other selection pressures at different mucosal sites. Future investigations of subtle differences in *tprK* diversity will require techniques that more confidently resolve rare alleles^30^. Additionally, we reveal increased *tprK* diversity in the SS14 lineage relative to Nichols and attribute it to an increased repertoire of unique donor sequences resulting from a 51 bp insertion in a V7-specific DC. As this increased diversity results in multiple SS14 lineage-specific antigens, enhanced *tprK* diversity may provide a fitness advantage to SS14 strains by enabling more variants to evade anti-*tprK* responses, potentially accounting for SS14’s expansion and prevalence globally^8^.

In conclusion, we find the use of inter-sample Bray’s dissimilarity scores of *tprK* V regions to be a promising tool to infer syphilis transmission. Future work involving wider sampling, including more known transmission events, will be necessary to confirm these findings and further refine methods. Critically, studies that connect *tprK* variation with host immune responses will be essential to define how AV shapes immune evasion by *Tp*.

## Acknowledgements

The authors are grateful for the work of the UW Long Read Sequencing Center.

## Supplementary Methods

### Ethics approvals and enrollment criteria (Brazil, Peru)

Enrollment of adults in Brazil was approved by the Brazil: National Research Ethics Commission protocol 56591822.9.0000.5243 and screening performed according to Brazilian National Guidelines^1^. Enrollment of adults in Peru was performed under the University of Southern California IRB HS-21-00353, approved by the ethical review board at Universidad Peruana Cayetano Heredia in Lima on December 3rd, 2019 (UPCH IRB protocol 103093). Enrollment criteria and screening of the Peruvian cohort have been previously described^2^.

### DNA Extraction and qPCR

In the Buenos Aires cohort, DNA from 300 µL whole blood samples or dry swabs suspended in 400 μL of PBS was extracted using QIAamp DNA Blood Mini Kit (Qiagen, Germany) according to the manufacturer’s instructions. Then, the samples were eluted in Qiagen AE buffer to a final volume of 200 µL for whole blood samples and 150 μL for swabs. Swabs from Brazil were collected in 2 mL of DNA/RNA shield stabilization solution and extracted using the Quick-DNA 96 Kit (Zymo Research; Irvine, CA). Peruvian swabs were collected in DNA lysis buffer (10 mM Tris-HCl pH 8.0, 0.1 mM EDTA, 0.5% SDS) and extracted using the QIAamp DNA Mini Kit. *T. pallidum* DNA was detected via semi-quantitative PCR for the single copy locus *tp0574* as previously described^2^. Samples were considered positive when *tp0574* cycle threshold was less than 40.

### T. pallidum WGS

Sequencing was attempted on samples with more than 25 *tp0574* copies/μL DNA eluate, equivalent to a minimum of approximately 200 genome copies into library preparation. Briefly, genomic DNA was fragmented using the KAPA HyperPlus kit (Roche) and TruSeq unique dual indices (Illumina) ligated prior to 14-16 cycles of pre-capture PCR. Hybrid capture using our custom DNA probes was performed using the manufacturer’s instructions (Integrated DNA Technologies) prior to 14-16 cycles of post-capture amplification. Paired-end 150nt reads were generated on Illumina instruments including the HiSeqX, Novaseq6000, and NextSeq2000.

### *T. pallidum* variant calling

Raw fastqs were newly generated or downloaded from the NCBI Sequence Read Archive. Reads were preprocessed by removal of human reads with Kraken2 v2.1.3^3^ and quality and adapter trimmed with Trimmomatic v0.39^4^, then filtered using BBDuk v39.10^5^ at low stringency against the SS14 reference genome (NC_021508) to identify reads containing *Tp* sequences. From these, *Tp* rRNA and tRNA were further filtered at high stringency, requiring 99% identity to the conserved *Tp* references. Filtered *Tp* reads were then mapped against the SS14 reference genome using bowtie2 v2.5.2^6^ with default parameters, deduplicated with Picard v2.26.3^7^ and variants called using GATK v4.6.1^8^ HaplotypeCaller assuming ploidy of 1 and maximum indel size of 130. Joint genotyping was performed using GATK and filtered with bcftools v1.2^9^ to a minimum depth of 3 reads and allele frequency >0.8. Masking of previously identified recombinant loci^10–12^ and regions with poor assembly quality, such as homopolymers, was performed prior to SNV-only phylogenetic inference.

### Custom *tprK* analyses

The number of alleles in each sample was defined as the number of unique full-length *tprK* sequences exceeding 0.2% abundance in the technical replicate of each sample with the highest sequencing quality score. K-mers of *tprK* alleles and donor cassette loci were generated using Jellyfish v1.2^13^. High-confidence *tprK* alleles used to evalutate intra-and inter-sample distance (**Figure 5** and **Figure S11**) are defined as those present in both technical replicates. Inter-sample Pearson correlations between V regions were calculated as previously described^14^. Inter-sample Bray’s distance was calculated for lengths of k=9, 11, 13, 15, 17, 19, 21, 23, 25, and 27 using V region-derived k-mers aggregated per sample (ie, without consideration for full-length *tprK* alleles). Inter-allele Bray’s distance was calculated using k-mers of lengths 9-27 for each pair of alleles and the median inter-allele distance for each sample pair used for statistics and visualization. Inter-sample V region and allele-derived Bray’s distances were summed across all values of k. Pairwise global sequence alignments were performed with mafft v7.41^15^ between individual V regions and inter-allele alignment distance evaluated as the sum of V1-V7. The median inter-allele alignment distance for each sample pair was used for statistics and visualization.

**Supplementary Table 2:**
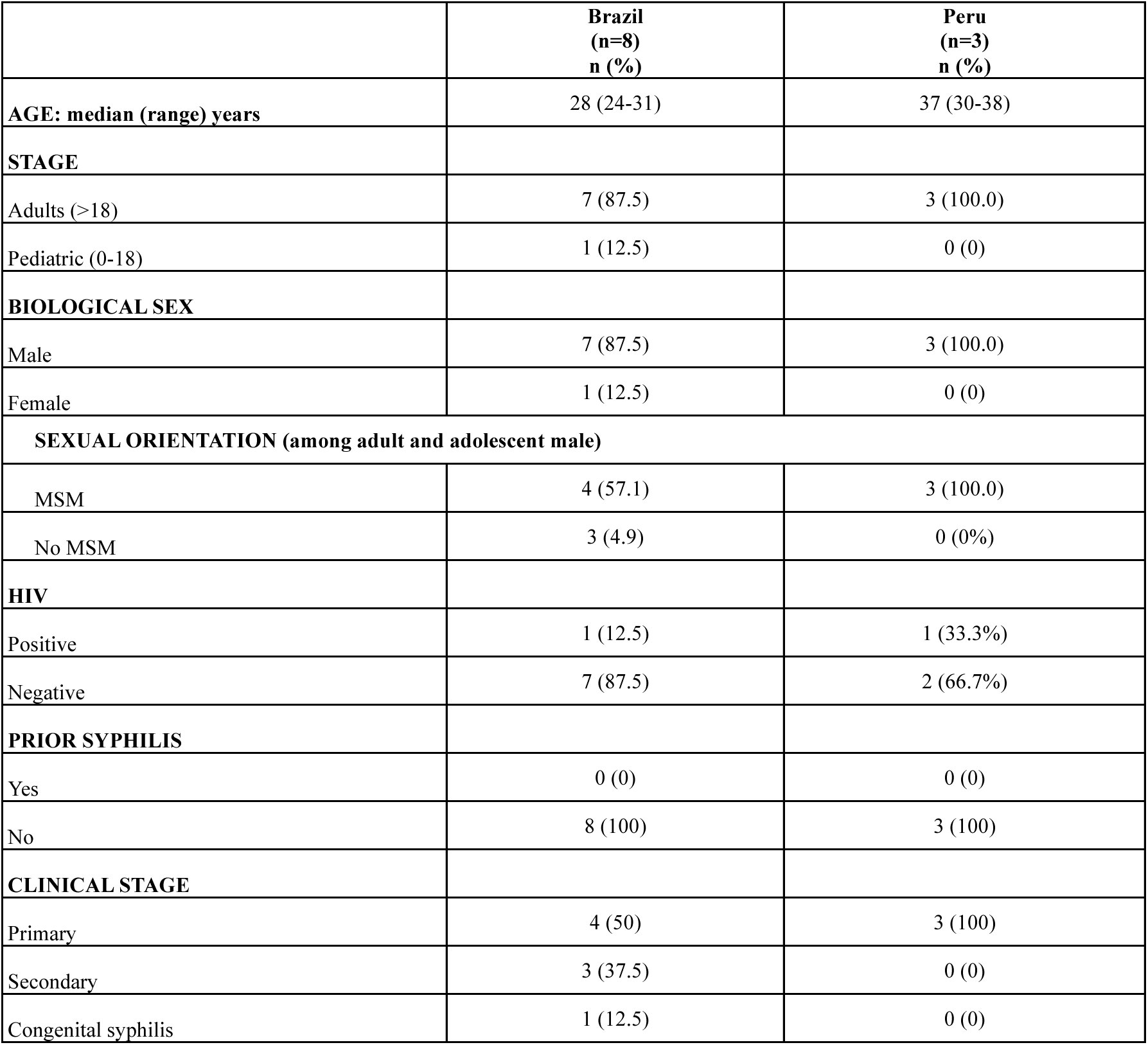
Demographic characteristics of participants from Brazil and Peru with WGS newly included in this study.

|  | **Brazil**<br>**(n=8)**<br>**n (%)** | **Peru**<br>**(n=3)**<br>**n (%)** |
| --- | --- | --- |
| **AGE: median (range) years** | 28 (24-31) | 37 (30-38) |
| **STAGE** |  |  |
| Adults (>18) | 7 (87.5) | 3 (100.0) |
| Pediatric (0-18) | 1 (12.5) | 0 (0) |
| **BIOLOGICAL SEX** |  |  |
| Male | 7 (87.5) | 3 (100.0) |
| Female | 1 (12.5) | 0 (0) |
| **SEXUAL ORIENTATION (among adult and adolescent male)** |  |  |
| MSM | 4 (57.1) | 3 (100.0) |
| No MSM | 3 (4.9) | 0 (0%) |
| **HIV** |  |  |
| Positive | 1 (12.5) | 1 (33.3%) |
| Negative | 7 (87.5) | 2 (66.7%) |
| **PRIOR SYPHILIS** |  |  |
| Yes | 0 (0) | 0 (0) |
| No | 8 (100) | 3 (100) |
| **CLINICAL STAGE** |  |  |
| Primary | 4 (50) | 3 (100) |
| Secondary | 3 (37.5) | 0 (0) |
| Congenital syphilis | 1 (12.5) | 0 (0) |

## Supplementary Analysis

### Extended clinical findings (Buenos Aires)

All primary syphilis cases in adults (18 male and 3 female), except one, had a chancre at sites of sexual exposure while two cases presented with Folman’s Balanitis (one patient had both a penis chancre and balanitis). Three primary syphilis cases had VDRL negative results, including one person living with HIV that presented with a genital chancre and darkfield positive result. Predominant symptoms among secondary syphilis cases (60 male and 32 female) were lymphadenopathy (65/92, 70.7%) and rash (52/92, 56.5%). Rash types included macular (16 cases), maculopapular (16), papular (15), and papulosquamous (5). Some patients also expressed oral or tongue plaque lesions (33/92, 35.9%), genital or perianal condyloma lata (30/92; 32.6%), and genital erosions (17/92, 18.5%). Five patients also had alopecia (5/92, 5.4%).

Congenital syphilis cases (one male and three females) were newborns and three had early signs of congenital syphilis including pemphigus-like lesions, hydrops fetalis, and hepatosplenomegaly. All primary syphilis cases were adolescents (two males and one female) with genital chancre. Secondary syphilis cases included 12 children (five males and seven females) and nine adolescents (five males and four females). Among them, common symptoms included macular or maculo-papular rash (7/21, 33.3%), genital or perianal condyloma (13/21, 61.9%) and palate erosion (5/21, 23.8%) patients. Few patients expressed lymphadenopathy (4/21, 19.0%), alopecia (2/21, 9.5%), and genital erosions (2/21, 9.5%).

### Lesion-specific SNVs in *tp0006*

We recovered near-complete genomes from multiple specimens from 20 individuals and found no evidence of intra-host evolution in 19 of the matched sets. In one sample pair, taken from perianal condyloma and skin rash from an individual (GSPAR095M) with secondary syphilis, two clonal SNVs (D136G in the perianal lesion and M273I in the axilla skin sample) were seen in *tp0006,* encoding an accessory protein of unknown function^16^. Analysis of the predicted structure using Foldseek^17^ revealed TP0006 to have structural homology to substrate binding domains of ATP-binding cassette transporters (**Figure S2;** Dr. Chad Brautigam, personal communication). Of note, the patient reported treatment with amoxicillin for an unrelated infection approximately six weeks prior to his presentation with secondary syphilis.

### Limited diversity in *tprK* constant regions

Among 79 samples from 54 patients with matched WGS, *tprK* constant regions showed limited sequence variation, with only six positions out of approximately 1,600 nucleotides different between Nichols and SS14 lineages and none in two previously identified long-lived T cell epitopes^18^. In samples from one patient (GSPAR031R), a 59-year-old cisgender heterosexual man newly diagnosed with HIV, *tprK* alleles from both genital mucosa and interdigital skin had a novel complex mutation resulting in conservative amino acid change T271A in the conserved region between V2 and V3, corroborated by WGS in four separate site samples from the same patient. A 13 nt sequence containing the altered sequence (**Figure S5A**) was identical to a motif within the conserved N terminus of *tpr* subfamily I, providing a donor sequence for a putative recombination event. Additionally, two patients had an A-to-C transversion that conferred a W516C mutation in the C-terminal tail of TprK (**Figure S5B**). In patient GSPAR093S the mutation was present in the resolving genital chancre at approximately 53% abundance and 90% in the oral lesion. In the matched WGS sample from the genital lesion, no other genomic positions were mixed, suggesting *de novo* mutation rather than co-infection. Using WGS data, the W516C variant is found between 10-100% allele frequency in 124/1,879 (6.6%) samples (**Figure S5C**).

---

**Figure S1:**
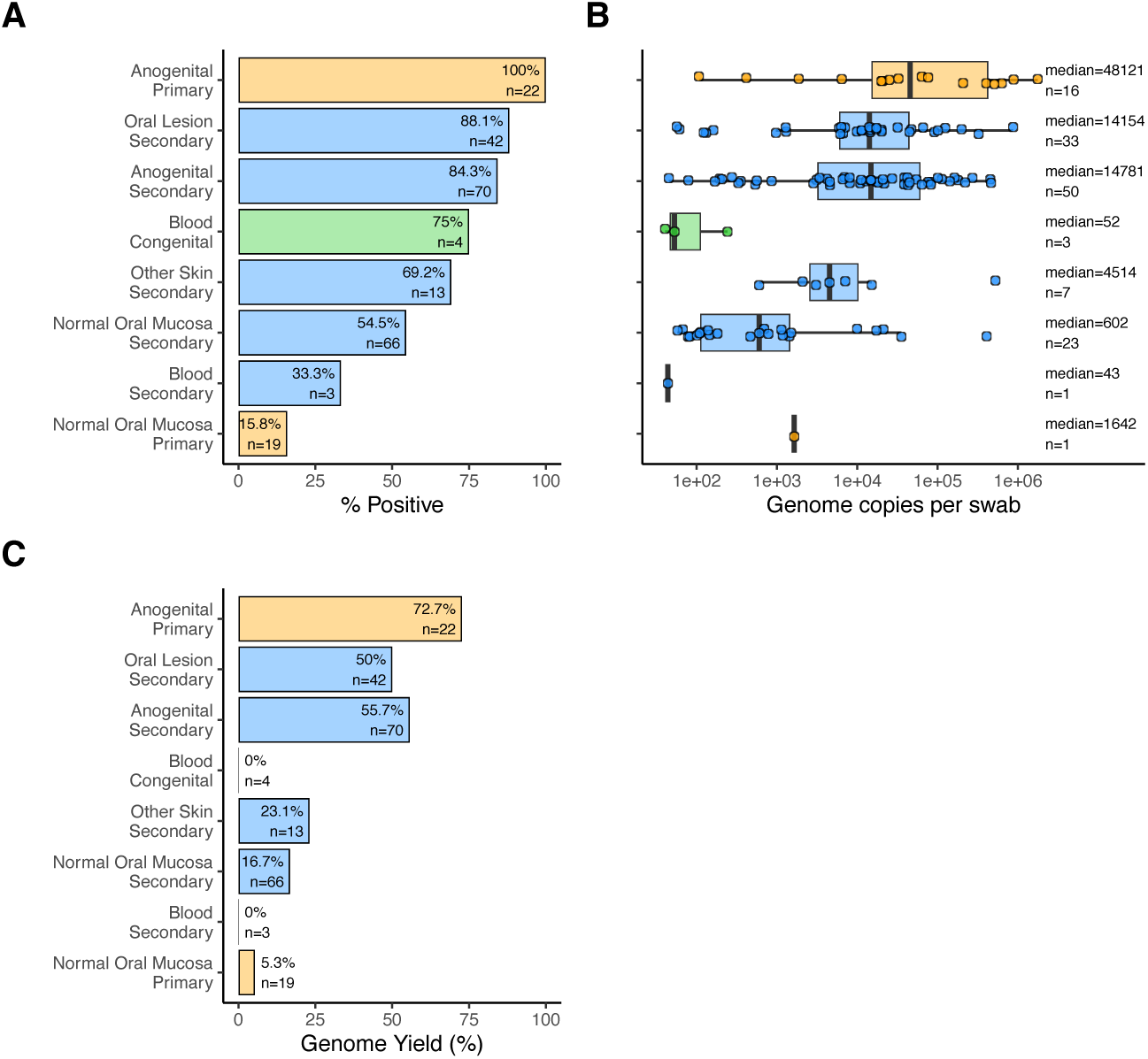
*T. pallidum* molecular positivity and treponemal burden by sample anatomic location and patient’s clinical stage. A) Percent of samples positive for the single copy locus *tp0574*. B) Extrapolated total *Tp* genome copies from a single swab or 300 μL whole blood, C) Percent of samples that yielded a complete genome

**Figure S2:**
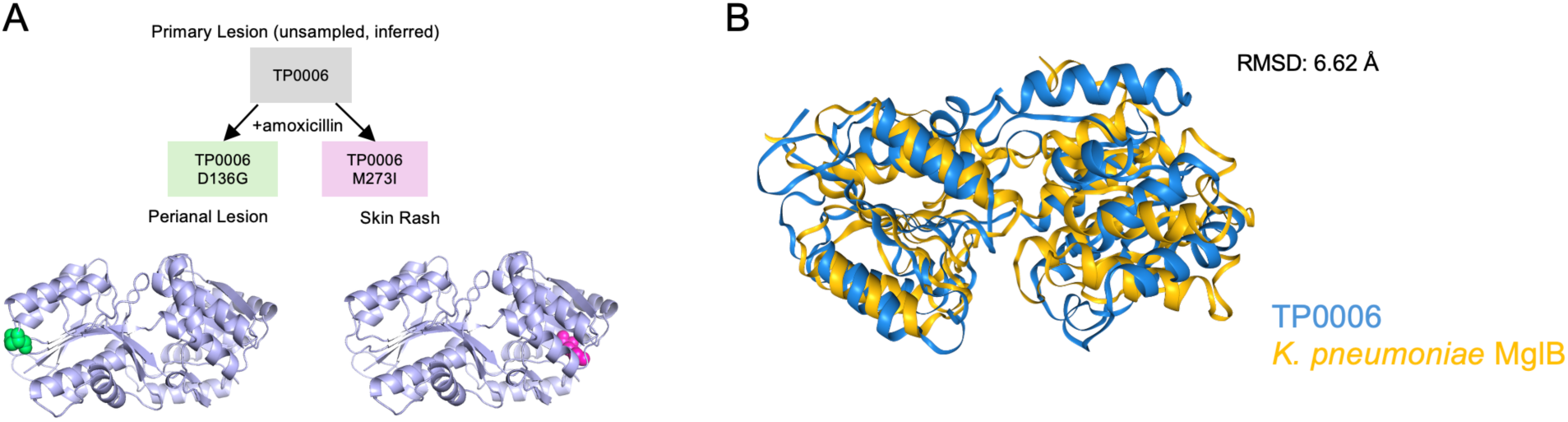
Mutations to *tp0006*, encoding a protein of unknown function. A) Two separate mutations were found in *T. pallidum* DNA recovered from a male patient’s perianal and skin lesions relative to the inferred genotype of the infecting strain, which was not sampled. Mutations are shown on the AlphaFold3-predicted structure of TP0006. B) TP0006 has structural homology to the substrate binding proteins of ABC transporters such as MglB. RMSD, root mean squared deviation.

**Figure S3:**
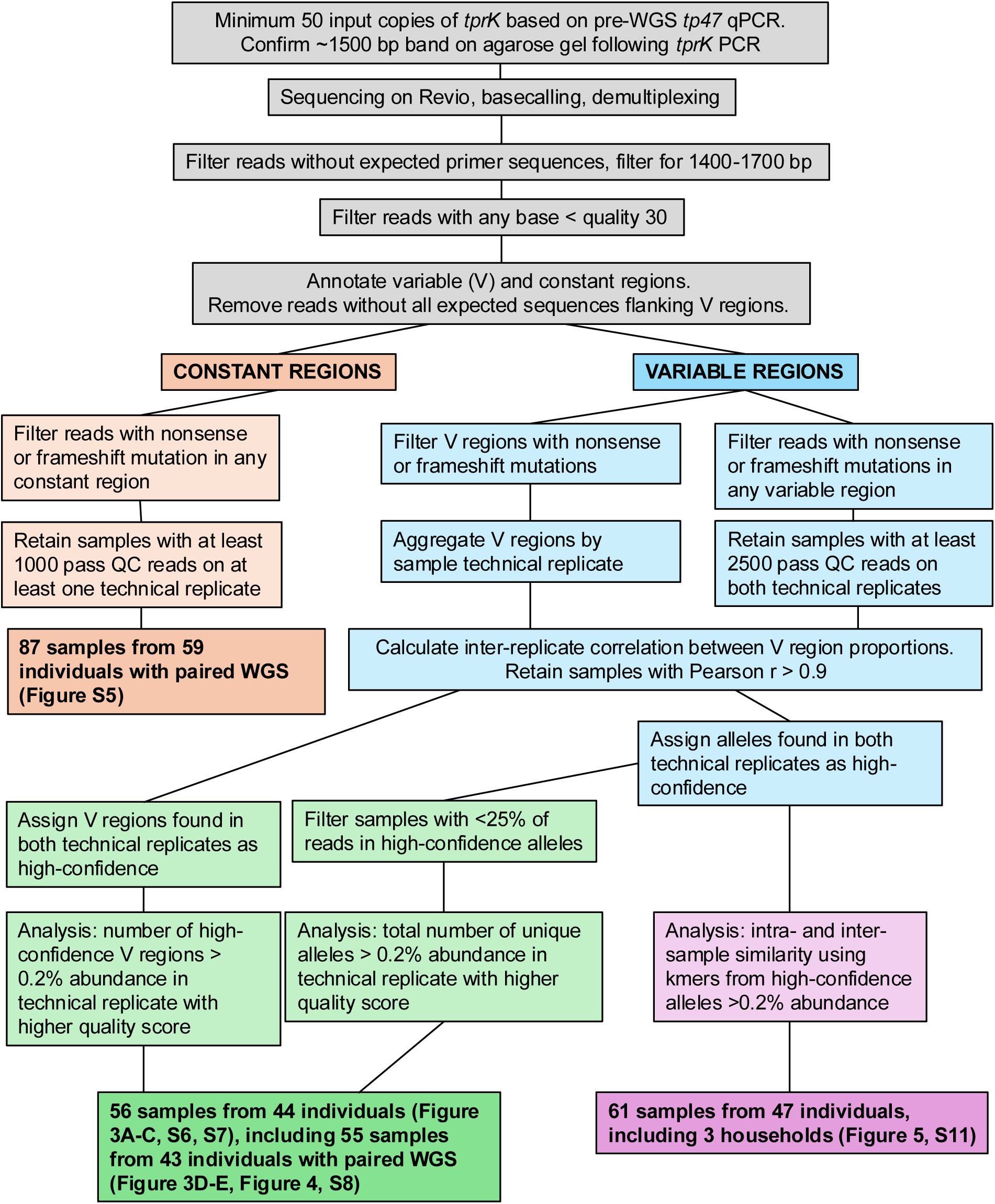
Quality control and filtering processes for each *tprK* analysis.

**Figure S4:**
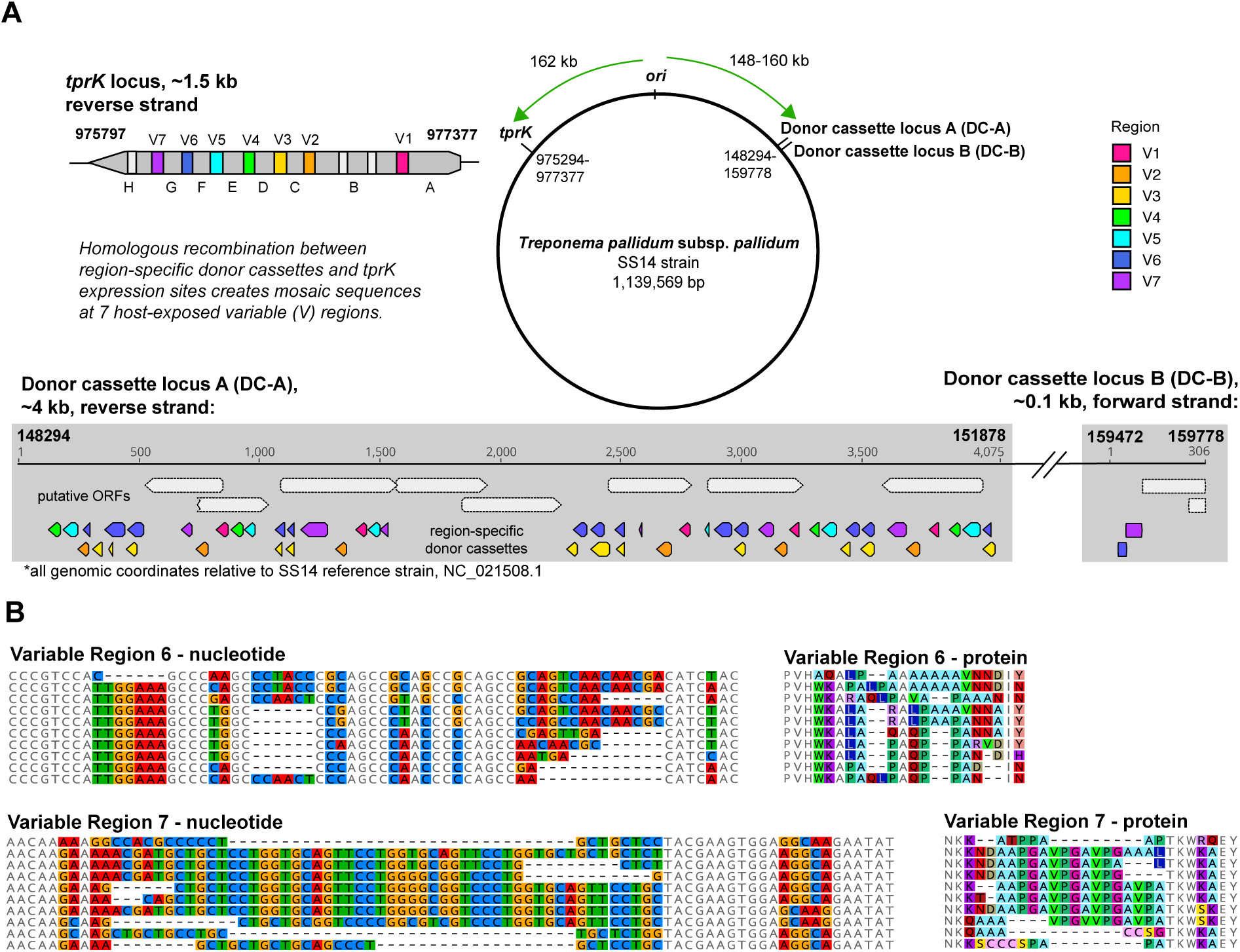
Gene conversion between donor cassettes and expression sites results in antigenic variation in seven variable loops of TprK. A) The *tprK* locus is found 162kb from the origin of replication on the reverse strand of the *T. pallidum* chromosome. *tprK* encodes a ∼500 residue predicted outer membrane protein. Seven of its ten variable loops undergo antigenic variation (colored). Constant regions A:H are found between variable regions. Two loci encoding donor cassettes (DCs) are 148 and 159 kb from the origin of replication, respectively. DC-A contains 51 discrete sites with donor cassettes on the reverse strand. DC-B contains 2 sites with donor cassettes on the forward strand. All DCs are V region-specific, with regions V3 (yellow) and V6 (blue) having overlap. DCs overlap with several transcribed open reading frames (ORFs). B) Examples of ten V6 and V7 sequences (out of 1020 and 313, respectively). Both nucleotide and amino acid variability are shown.

**Figure S5:**
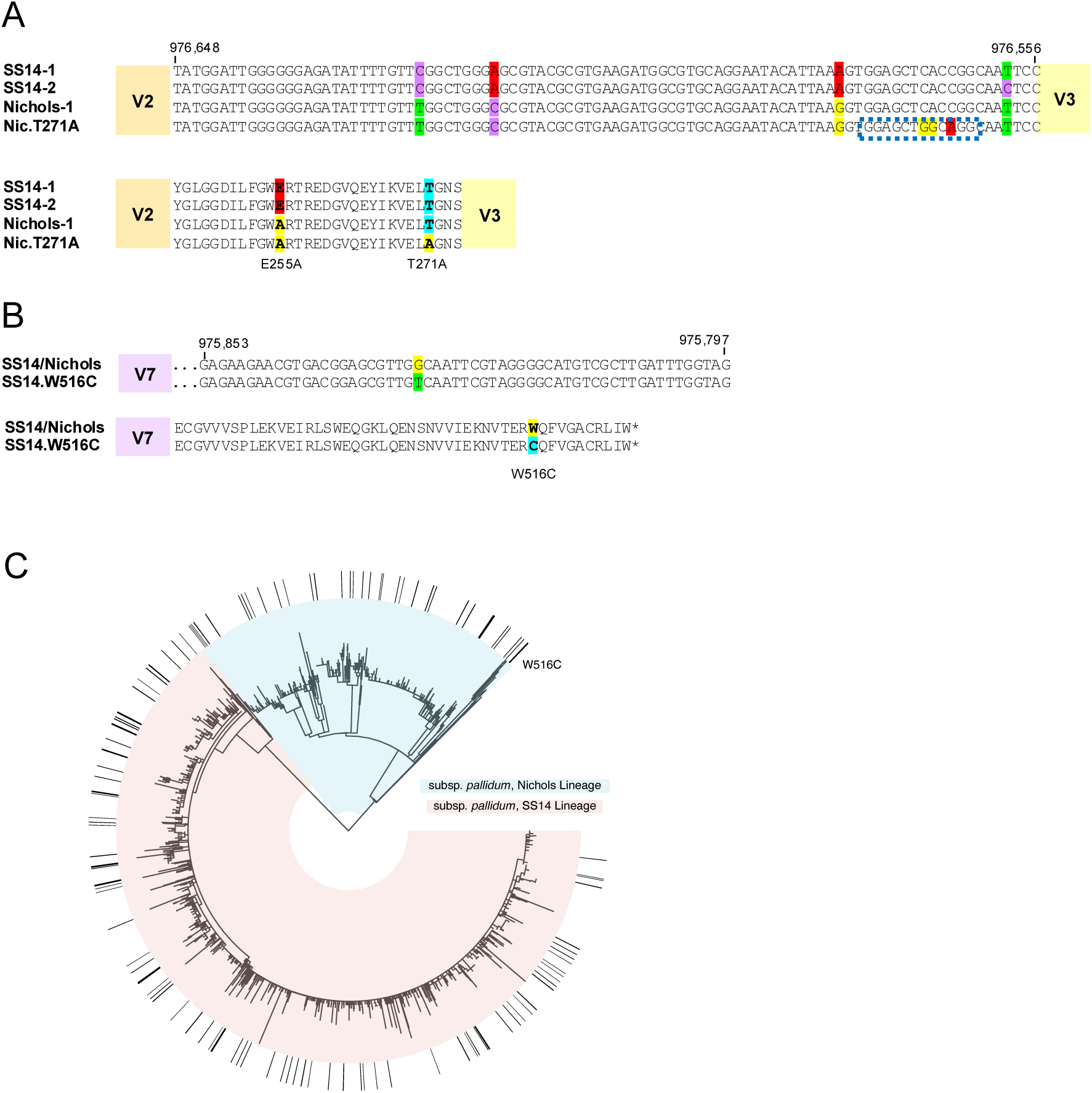
Nucleotide and amino acid differences found in *tprK* constant regions. A) Lineage-defining mutations between V2 and V3. In one patient, a recombination event from a donor site in a subfamily I *tpr* (blue box) was found in samples from four unique body sites, resulting in T271A. B) A SNV resulting in W516C is fully fixed in one patient and incompletely fixed in another. C) Global phylogeny of 1879 strains from Figure 1A. Nichols-lineage samples have a blue background and SS14 red. The outer ring denotes strains with W516C fully or partially fixed.

**Figure S6:**
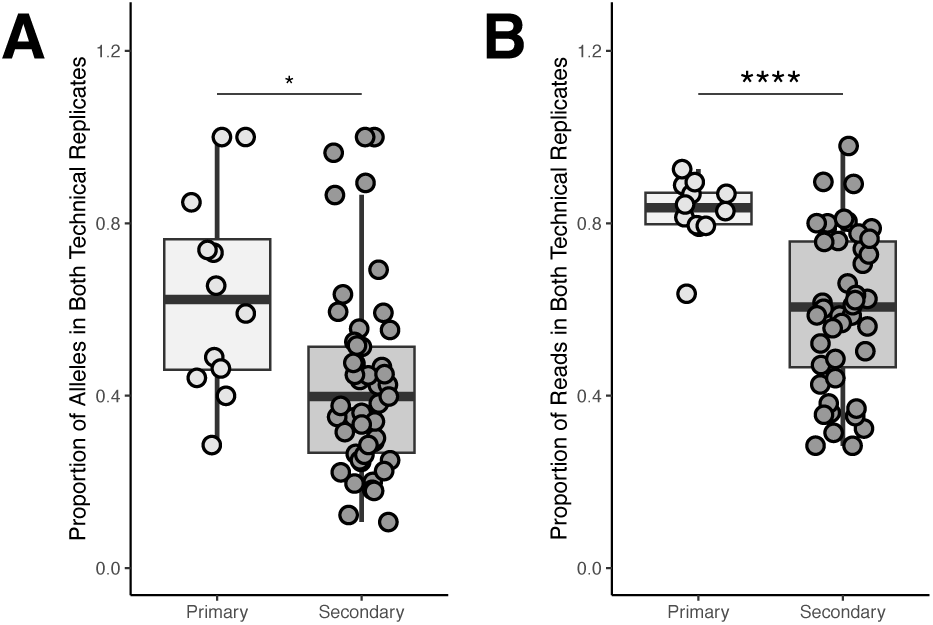
The proportion of unique *tprK* alleles (A) and the percent of total reads (B) in high confidence alleles varies by stage. T-test, *p<0.05, ****p<0.0001

**Figure S7:**
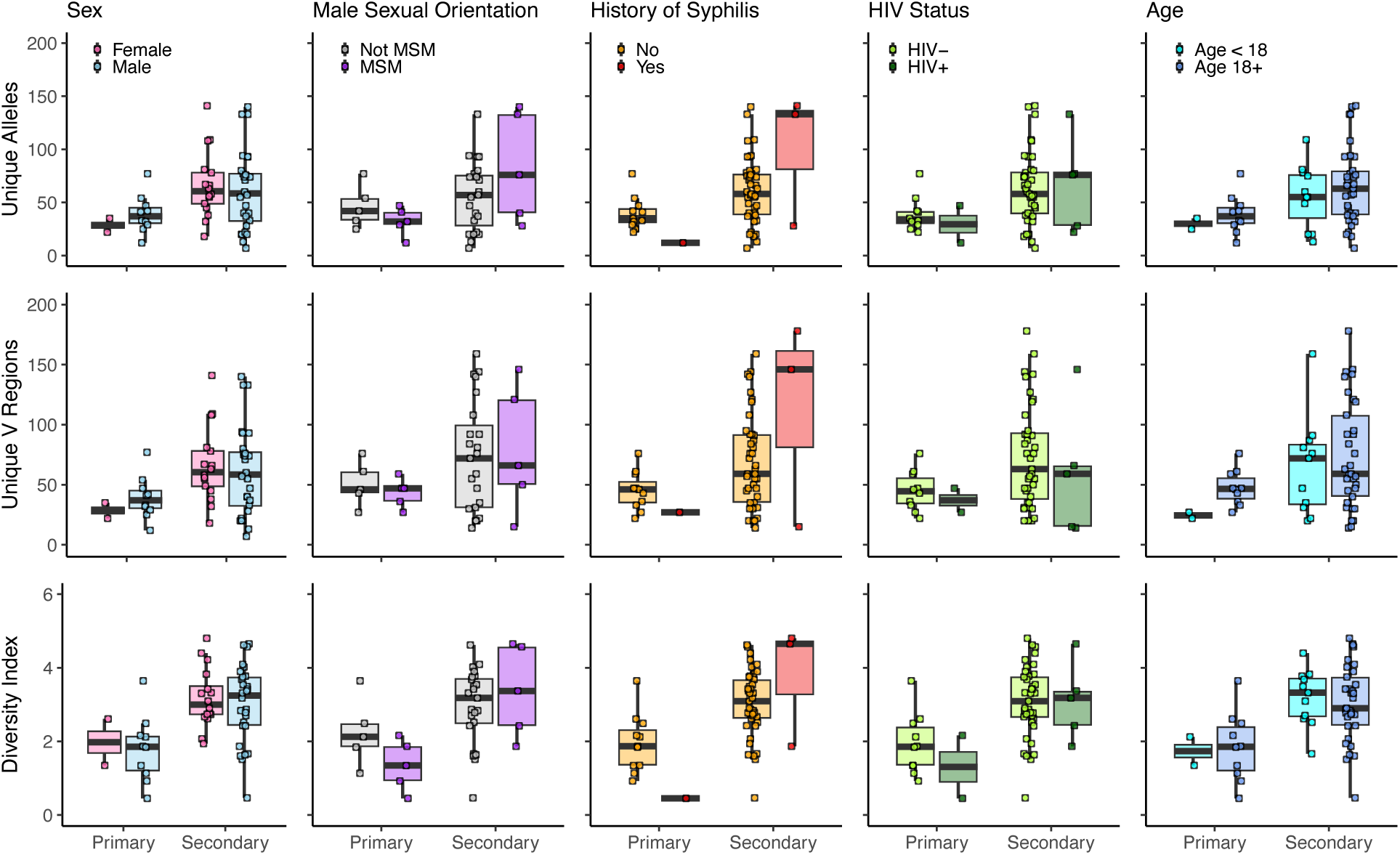
No differences were seen in the number of *tprK* alleles, V regions, or diversity index by patient sex, sexual orientation, reinfection, HIV status, or age.

**Figure S8:**
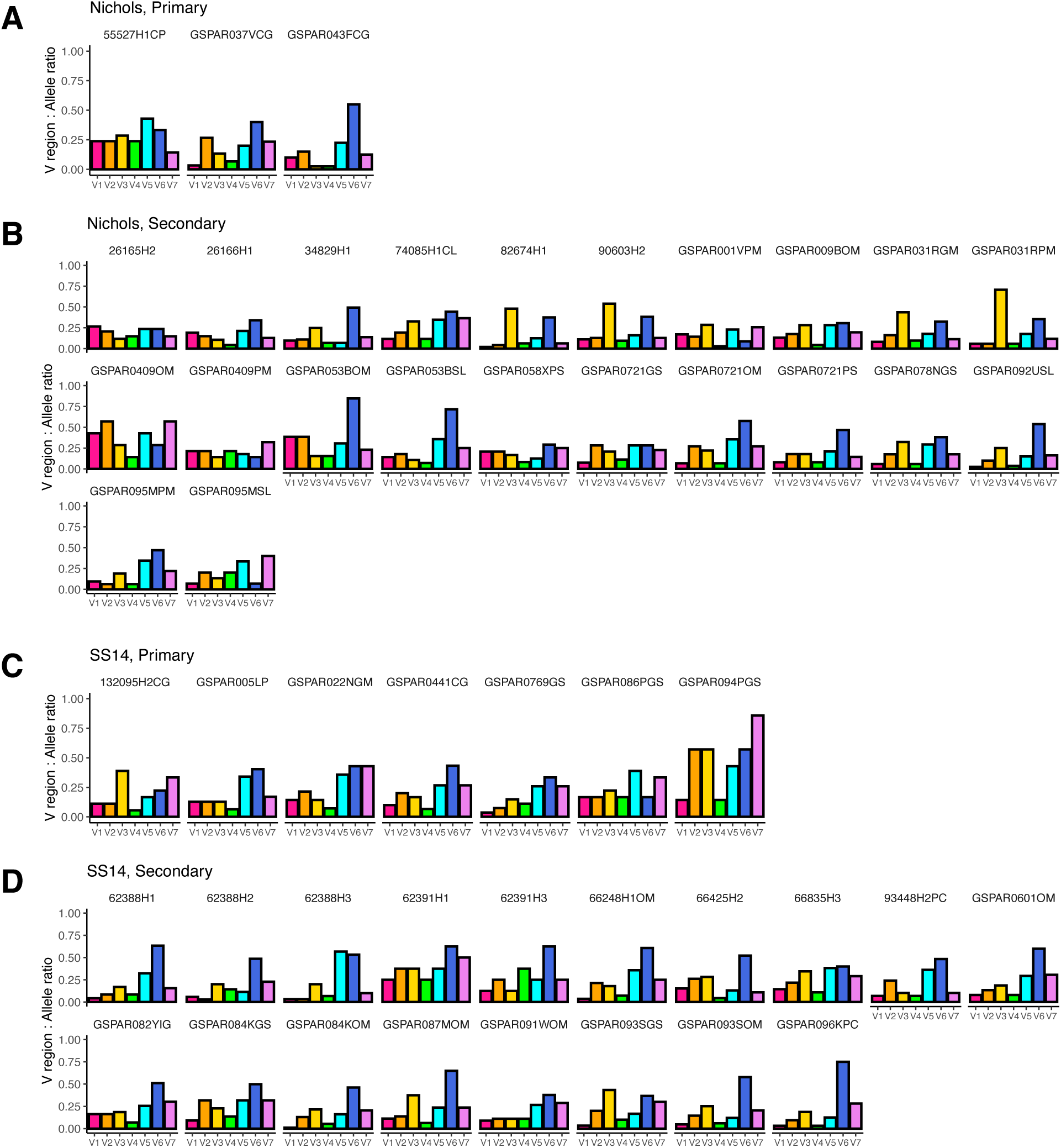
Distribution of unique V regions. The proportion of unique V regions to the number of unique *tprK* alleles is shown for Nichols lineage (A, B) and SS14 lineage (C, D) from primary and secondary syphilis cases. For example, a sample with 20 unique *tprK* alleles, 2 unique V1 sequences, and 10 unique V6 sequences would have a V1 V:Allele ratio of 0.1 and a V6 V:Allele ratio of 0.5.

**Figure S9:**
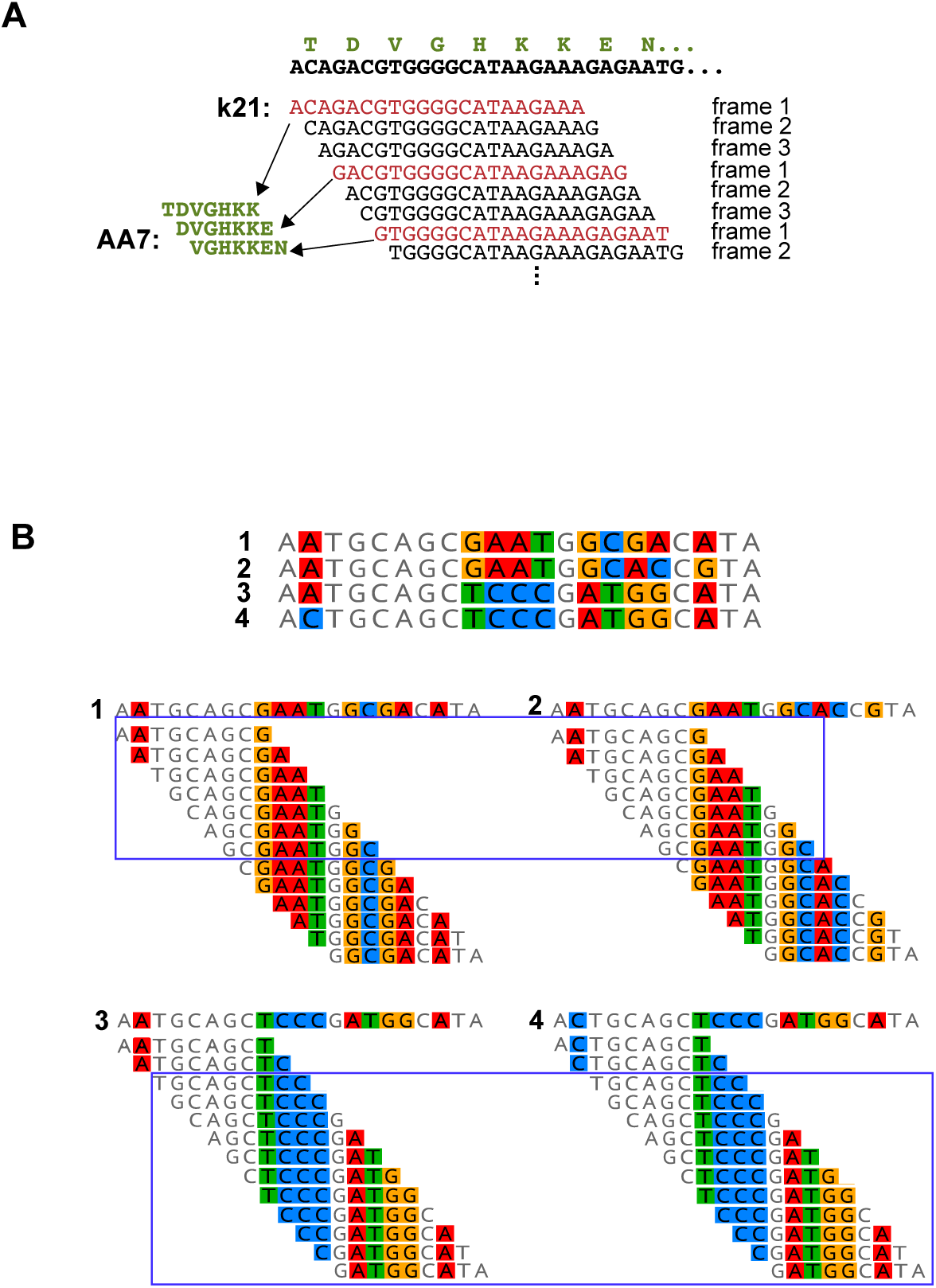
*tprK* analysis using kmers. A) Each *tprK* sequence fragment is broken into overlapping sequences of length k (k=21 is shown). Kmers in frame 1 encode amino acid fragments (AA-mers) of length k/3. B) Similarity between *tprK* alleles can be evaluated using the number of shared kmers/AA-mers. Although all four sequences shown derive from V4, those in the top row share no kmers with the bottom row.

**Figure S10:**
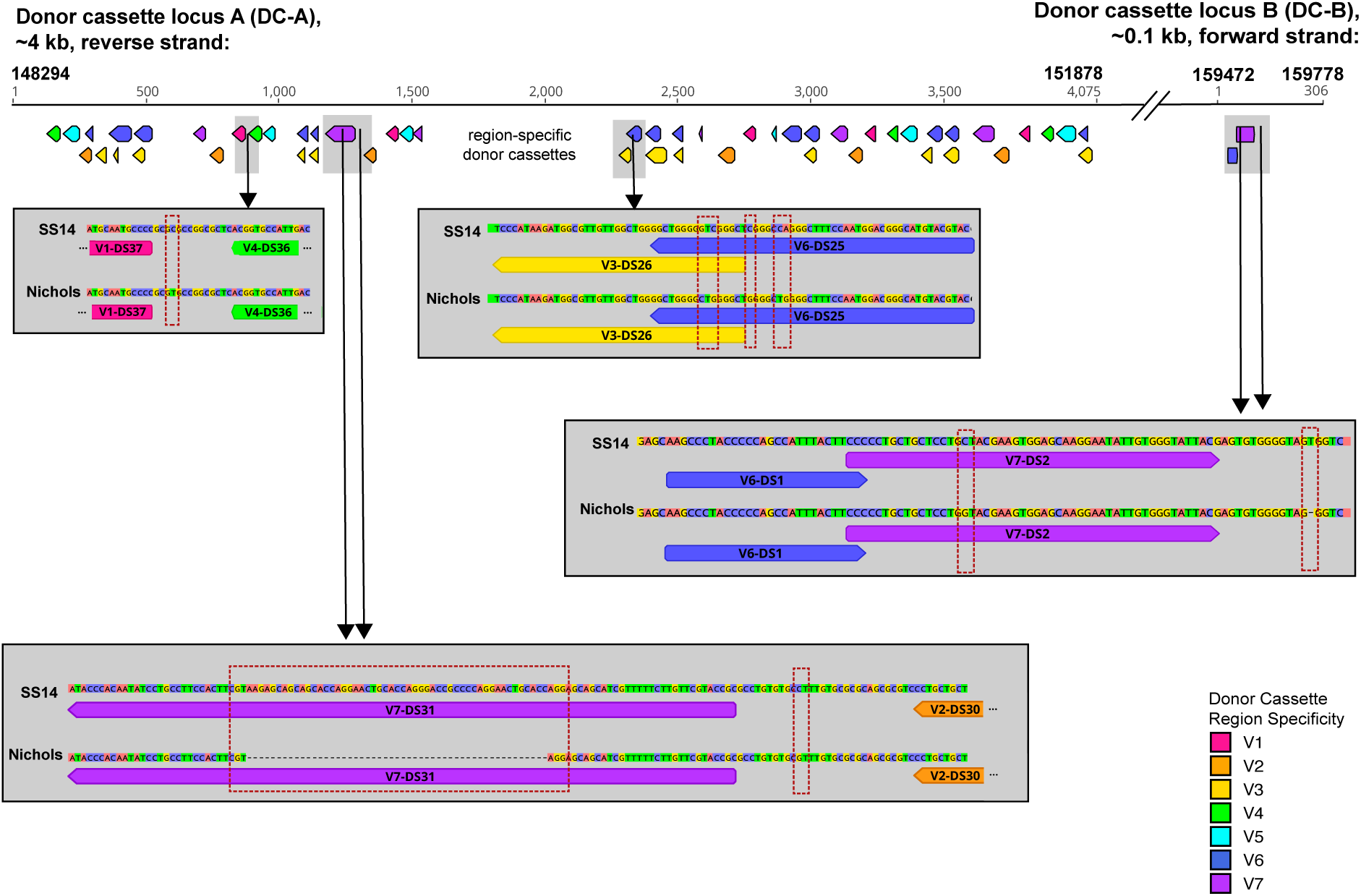
Differences in donor cassette loci between the Nichols and SS14 lineages (grey background) are confined to donor cassettes in regions V3-DC25/V6-DC26, V7-DC31, and V7-DC2, as well as sequences between cassettes.

**Figure S11:**
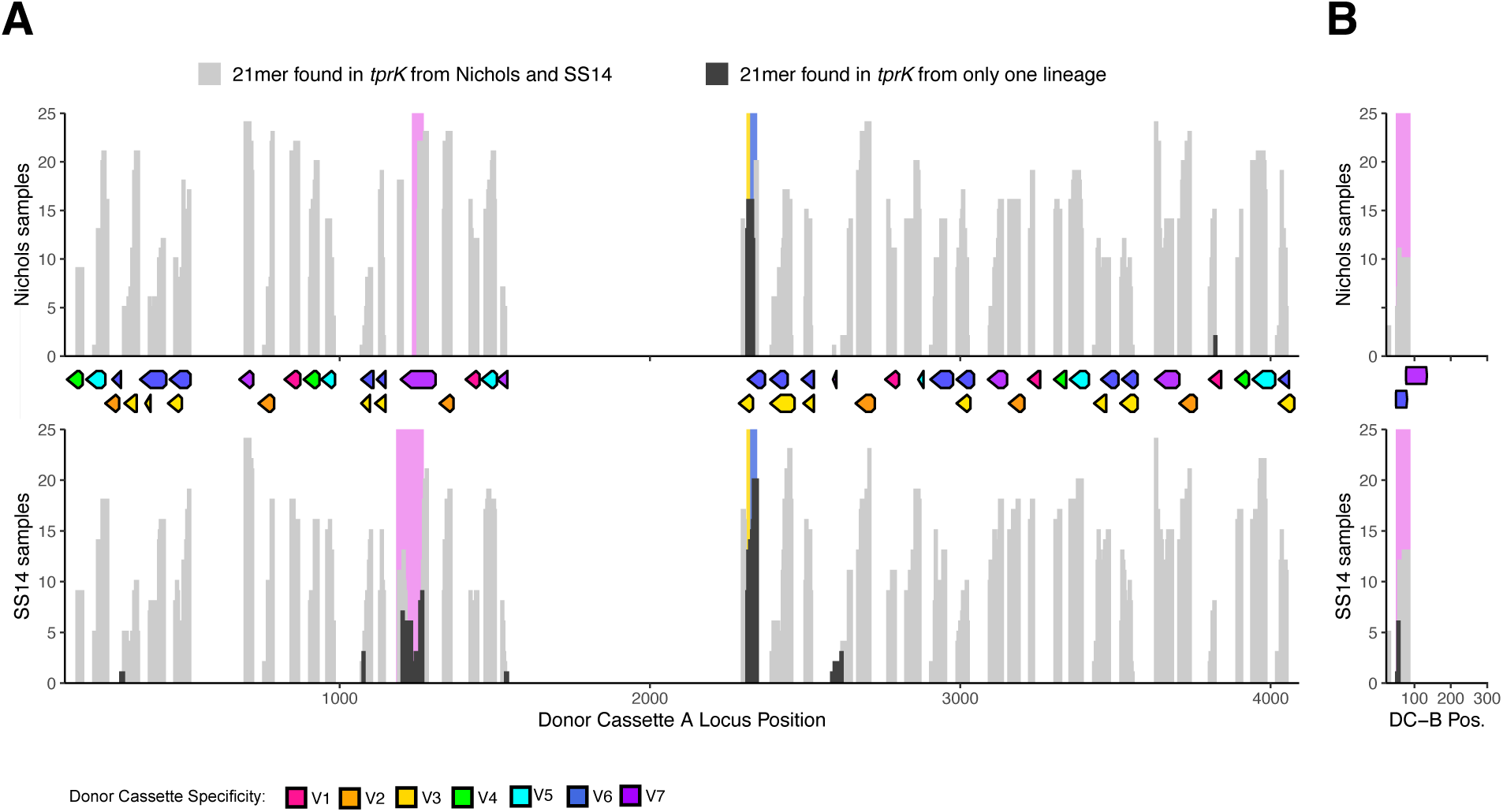
Lineage-specific DC-derived sequence donor location in Nichols and SS14. Histogram of the number of Nichols (top) or SS14 (bottom) samples containing at least one *tprK* allele using DC-derived 21-mer from each donor site in A) DC-A and B) DC-B. Donor cassettes colored by V region specificity in their relative positions are shown between the plots. DC-derived k-mers found in *tprK* sequences from both Nichols and SS14 lineages are shown in light grey. DC-derived k-mers found in only a single lineage are shown in dark grey. Sequences with differences between Nichols and SS14 that are within DCs are highlighted on the plot colored by V region specificity.

**Figure S12:**
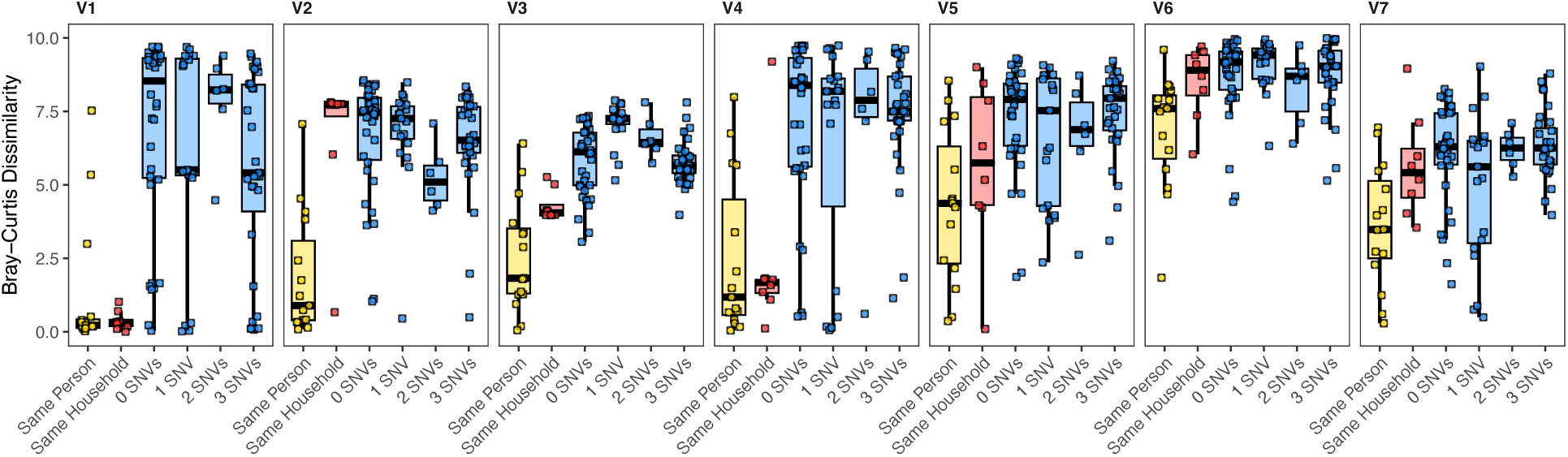
Inter-sample dissimilarity between *tprK* V regions. Dissimilarity of k-mers of length 21 at each sample relationship type varies by V region, with V1 and V4 generally similar within a single patient or household, while V6 is highly variable

